# Linking MSMEG_1353 to lipid metabolism and envelope integrity in *Mycolicibacterium smegmatis*

**DOI:** 10.64898/2025.12.18.695065

**Authors:** Ziwen Xie, Trillion Surya Lioe, Abhipsa Sahu, Jiahao Cui, David Ruiz-Carrillo, Tatsuhiko Kadowaki, Boris Tefsen

**Affiliations:** Department of Biosciences and Bioinformatics, Xi’an Jiaotong-Liverpool University, Suzhou, Jiangsu, China; Biophysical Structural Chemistry, Leiden Institute of Chemistry, Leiden University, Einsteinweg 2333 CC, Leiden, Netherlands; International Centre for Genetic Engineering and Biotechnology, Taizhou, Jiangsu, China; European Molecular Biology Laboratory, Notkestraße 85, 22607, Hamburg, Germany; Department of Molecular Microbiology, Utrecht University, Padualaan 8, Utrecht, Netherlands

**Keywords:** *Mycolicibacterium smegmatis*, *Rv0647c*, *MSMEG_1353*, mycolic acid synthesis, isocitrate lyase, glyoxylate

## Abstract

*Mycobacterium tuberculosis* (*Mtb*) poses a significant global health burden. Rv0647c, an essential Mtb cell wall protein, is a potential drug target. We studied its *Mycolicibacterium smegmatis* (*MSMEG*) homologue, MSMEG_1353, using a CRISPRi conditional knockdown. MSMEG_1353 depletion increased cell width and volume, delayed log-phase initiation, slowed aggregation, reduced biofilm formation, and heightened susceptibility to antibiotics and sodium dodecyl sulfate (SDS). AlphaFold, sequence alignment, and UNIPROT analyses suggest MSMEG_1353 functions as a protein kinase, with conserved residues in intermediate high-confidence regions potentially forming an ATP-binding site. Investigating its role in cell envelope biosynthesis, mass spectrometry revealed elevated levels of mycolic acid biosynthesis proteins upon MSMEG_1353 knockdown. RT-qPCR confirmed upregulation of the fabD-acpM-kasA-KasB-accD6 operon, encoding key mycolic acid synthesis enzymes. Notably, a strong negative correlation with MSMEG_0911, the predominant isocitrate lyase, important in the glyoxylate cycle, was observed via both MS and RT-qPCR. Collectively, MSMEG_1353 deficiency compromises cell wall integrity, likely due to altered lipid composition resulting from dysregulated mycolic acid biosynthesis and lipid metabolism. These findings support the development of models explaining MSMEG_1353’s involvement in these pathways.

## 1. Introduction

*Mtb* causes tuberculosis (TB), and the morbidity and mortality of this disease can be exacerbated by other diseases and the development of multi-drug resistance, resulting in it being ranked 13th among the top 20 causes of human deaths (Bagcchi, 2023). DR-TB makes for an even more serious public health threat, and there are already more than 0.4 million estimated cases per year since 2015. Most potential drug targets are found in the cell envelope biosynthesis pathway and metabolism pathway of *Mtb* in recent years, suggesting those parts have high potential to explore new drug targets. The cell envelope of *Mtb* can change in thickness according to its different life stages and processes host immune modulatory functions (Dulberger *et al*., 2020). When considering the *Mtb* cell envelope from inside to outside, it is composed of the cytoplasmic membrane, peptidoglycan, arabinogalactan, mycolic acid, surface lipids and a capsule, and various proteins are found throughout the whole cell envelope as they are required for biosynthesis of the *Mtb* cell envelope, they contribute to the high impermeability of *Mtb*. Examples are kinases, transporters, regulators of cell wall synthesis and periplasmic cell wall assembly proteins (Dulberger *et al*., 2020, Chiaradia *et al*., 2017). Mycolic acid is one of the non-protein virulence factors which requires acetyl-CoA as precursor for biosynthesis. In *Mtb*, mycolic acids are constructed of 2-alkyl, 3-hydroxy fatty acids, the longest carbon-chains in nature, which is C_60_–C_90_ (Teramoto *et al*., 2015). In 2-alkyl, 3-hydroxy fatty acids, the α-chain is the saturated lipid at the 2 position and the meromycolate is the longer lipid on the 3-hydroxyl. Both the α-chain and the initial part of meromycolates are synthesised by fatty acid synthase (FAS) I enzymes, and FAS-II enzymes subsequently elongate the remaining part of meromycolates. After FAS-II synthesis, DesA, CmaA and MmaA protein families modify the meromycolates into six different types: α *cis/cis*, keto *trans*, keto *cis*, methoxy *trans*, methoxy *cis* and hydroxy *trans*. The α-chain and modified meromycolates are combined to form mycolic acid via polyketide synthase 13 (Marrakchi *et al*., 2014). Mycolic acid can be esterified with arabinogalactan. Others remain free, or are esterified with trehalose to form trehalose monomycolate and trehalose dimycolate. Several Serine/Threonine Protein Kinases (STPKs) can regulate the mycolic acid synthesis. STPKs can phosphorylate FAS-II enzymes, especially during dormant and latent phases of infection (Dulberger *et al*., 2020). *Mtb* is able to control mycolic acid both spatially and temporally to modify the immune response of the host during various infection stages. Proteins associated with mycolic acid biosynthesis and biosynthesis regulation are potential drug targets of *Mtb*.

The essentiality of the *Rv0647c* gene was demonstrated in a comprehensive screening of *Mtb* using Himar1 transposon mutagenesis (DeJesus *et al*., 2017). Previous studies have identified the presence of Rv0647c in both the cytosol and cell wall fraction of *Mtb H37Rv* through advanced techniques such as 2D liquid chromatography-mass spectrometry (LC-MS) (Mawuenyega *et al*., 2005) and nanoLC-MS/MS analysis, specifically in the membrane fraction (Xiong *et al*., 2005). Furthermore, MS has confirmed its presence in both the membrane protein fraction and whole cell lysates of *Mtb H37Rv*, while it was not detected in the culture filtrate (de Souza *et al*., 2011). The Rv0647c shows a conserved neighbouring with gene of lipG, a phospholipase/thioesterase involved in cell envelope synthesis (Santucci *et al*., 2018). Moreover, a STRING analysis (Szklarczyk *et al*., 2019) shows Rv0647c is in a cluster of mycolic acid cyclopropane synthase proteins (mmaA1, lipG, Rv0648c and fabD2). Furthermore, *Rv0647c* was found to be under control of the *SenX3 | Rv0490-RegX3 | Rv0491* regulon (Parish, 2003), which responds to low inorganic phosphate and nutrient starvation. When *SenX3* was deleted, the expression level of *Rv0647c* decreased corresponding with the expression level of several genes potentially involved in cell envelope biosynthesis, like *Rv1252c (lprE*) which codes for a lipoprotein and *Rv1518* which is involved in exopolysaccharide synthesis (Parish, 2003). These results combined indicate that Rv0647c is an essential protein which may play a crucial role in cell envelope biosynthesis regulation. The *Rv0647c* homologue in *Mycolicibacterium smegmatis* (*MSMEG*) has 84% identity (see Fig. S1 for alignment of MSMEG_1353 and Rv0647c) and is essential according to a pooled CRISPRi library (de Wet *et al*., 2018).

Here, we created *MSMEG_1353* conditional knockdown and overexpression strains and used those to address its function with a variety of approaches. Diminished amounts of MSMEG_1353 in the bacteria resulted in numerous phenotypes that all point to an altered and weakened cell envelope. Expression analysis at protein and mRNA level indicated that knockdown of MSMEG_1353 in inversely correlated with the expression of MSMEG_0911 (ICL), an isocitrate lyase that is part of the glyoxylate cycle, as well as with mycolic acid biosynthesis genes (*MSMEG_4325* (*fabD*) and *MSMEG_4327-4329* (*kasA, kasB, accD6*)). Together, these findings strongly suggest that MSMEG_1353 plays a role in maintaining cell wall integrity and let us propose a model in which it acts as a regulator in cell wall synthesis.

## 2. Material and Methods

### 2.1 Bacterial strains and standard growth conditions

All bacterial strains were documented in Table S1. Unless otherwise specified, all the reagents were purchased from Sigma-Aldrich, China. *Escherichia coli* (DH5α) was grown in LB medium or LB agar plate for genetic manipulation. *MSMEG* strains were grown in 7H9 medium (Difco) or 7H10 apar plate supplemented with 10% (v/v) albumin dextrose saline (ADS) (10 g of albumin, 4 g of D (+) - glucose and 1.7 g of sodium chloride in 200 mL of ultra-pure water), 0.5% glycerol (v/v), 0.05% Tween-80 (v/v) (Solarbio, China) (Gu *et al*., 2015) and appropriate antibiotics as mentioned in Table S1. All bacterial strains were grown overnight at 37 °C, shaking at 200 rpm.

### 2.2 Activation of conditional mutant strains

The ftsZi, 1353i, WT2P or WT strains were cultured by standard condition for around 18 hours. They were diluted to OD_600_ of 0.005 using fresh culture medium, and 200 ng/mL anhydrotetracycline (aTc) was added after 6 hours culture at 37 °C, shaking at 200 rpm. After another 18 hours of culture, these strains were used for experiments, and named as aTc-induced ftsZi, aTc-induced 1353i and aTc-induced WT2P strains and aTc-treated WT strain.

### 2.3 Overexpression plasmid construction

Genomic DNA was isolated from mc^2^155 using a standard protocol (Hu *et al*., 2021). *MSMEG* genomic DNA was used as the template for *MSMEG_1353* gene amplification with 1353-Forward-BamHI and 1353-Reverse-HindIII primers (Table 1) with a standard 50 μL PCR reaction mixture based on NEB Q5-high GC manufacturer’s instructions. PCR products were obtained by gel electrophoresis followed by gel extraction with standard protocol. Following the manufacturer’s instructions of restriction enzymes from NEB, the purified *MSMEG_1353* gene or pMV261 vector were digested by BamHI (NEB) and HindIII (NEB). Purified digested *MSMEG_1353* gene and pMV261 were mixed by 3:1 molar and ligated viaT4 DNA ligase (NEB) following its manufacturer’s instructions. All ligation mix was transformed into competent *Escherichia coli* (DH5α) with a standard protocol for constructed plasmid sequencing and amplification.

**Table 1.**
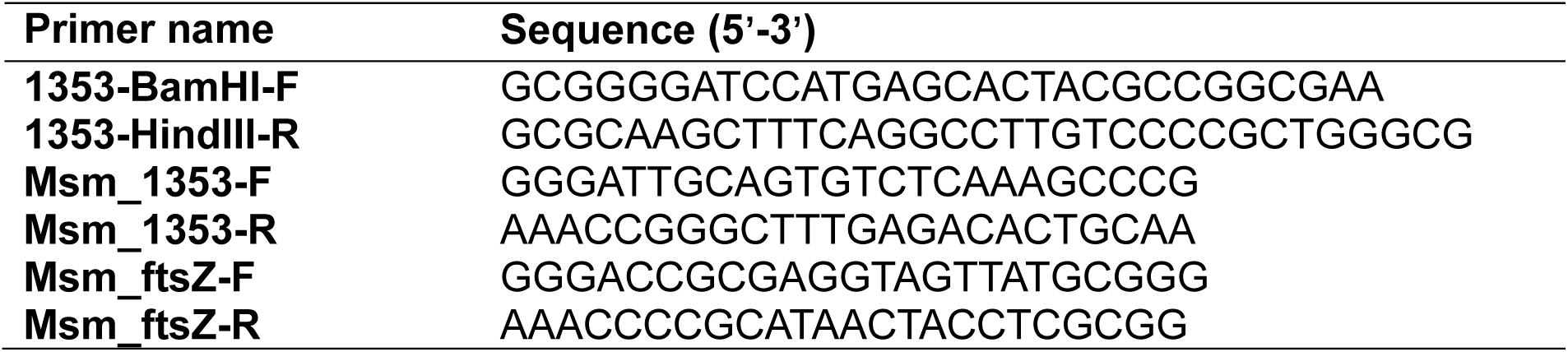
P rimers used for cloning.

### 2.4 CRISPRi plasmid construction

The sequences of the sgRNA oligos (Table 1) for making sgRNA anneals to the target gene and enables dCas9 to bind to it and block gene expression was obtained followed from the previously reported method without any change (Singh *et al*., 2016) and supported by the TEFOR website with the lowest off-target number and the highest outcome (http://crispor.tefor.net/crispor.py) (Concordet and Haeussler, 2018). The annealed oligos were inserted into pRH2521 plasmid for further transformation as reported method mentioned (Singh *et al*., 2016) and obtained constructed plasmids pRH2521-ftsZ and pRH2521-1353.

### 2.5 *MSMEG* strains construction

The method for *MSMEG* competent cell preparation was followed from a previous study (Hu *et al*., 2021). And constructed or empty plasmids were transformed into *MSMEG* mc^2^155 using the standard electroporation method (Gu *et al*., 2015).

### 2.6 mRNA isolation and RT-qPCR

Cells from the different *MSMEG* strains were obtained during logarithmic phase (OD_600_ between 0.8 and 1.2) and centrifuged at 8000 × *g* for 5 min and discarded the supernatant. The pellet was resuspended in 1 mL TRI reagent and mixed with around 50 μL 5 mm sized glass beads (Olan glass beads) in a mini bead-beater (Biospec) twice for 1 min on and 1 min off. Then heated at 60 °C 10 min and spin down at 12000 × *g* at 4 °C for 10 min. Then, the mRNA was extracted from supernatant by using the Innuprep Micro RNA kit (Analytikjena), following the manufacturer’s instructions. The cDNA was reverse transcribed from extracted mRNA by GoScript™ Reverse Transcription System (Promega), following the manufacturer’s instructions.

RT-qPCR was done with GoTaq® qPCR Master Mix (Promega) and primer sequences listed in Table 2. Reaction mix was made following the manufacturer’s instructions. The RT-qPCR was run in QuantStudio™ 5 System (Thermo Fisher Scientific). The *rpoD* (*sigA*) was used as the reference gene (Bustin *et al*., 2009) for cycle threshold (ct) value calculation via 2^−ΔΔCt^ method (Livak and Schmittgen, 2001).

**Table 2.**
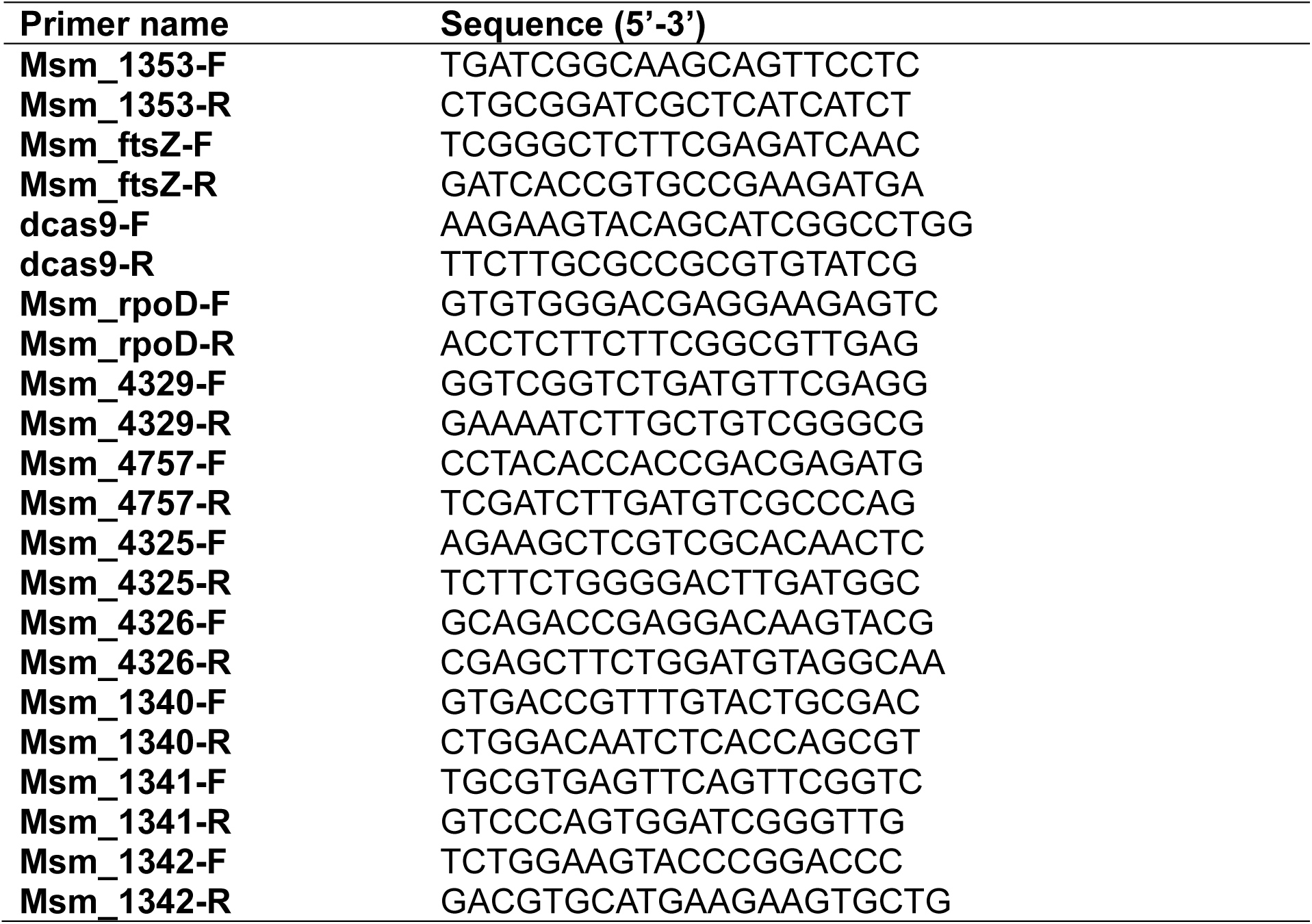
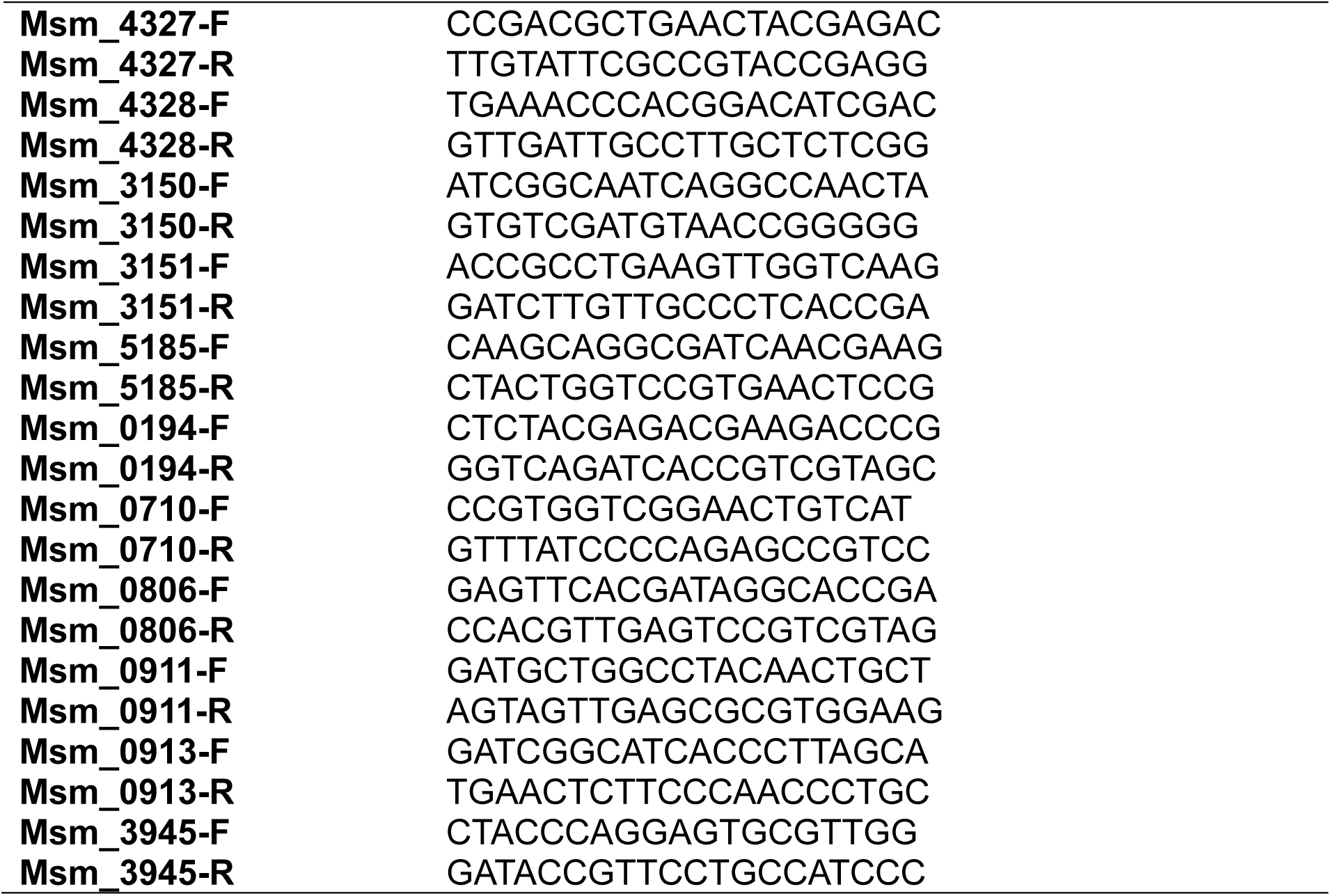
P rimers used in RT-qPCR.

### 2.7 Growth characteristics analysis

All strains were cultured started from an OD_600_ of 0.005 bacteria in 80 mL culture medium at 37 °C for 60 hours, OD_600_ value of each strain was measured every 3 hours by a spectrophotometer (BioPhotometer plus, Eppendorf). The growth curve of each strain was calculated in the nonlinear (curve fit) method using an exponential growth equation fit in OriginPro 2020 (OriginLab Corporation).

### 2.8 Light-microscopy and scanning electron microscopy (SEM)

For light-microscope observation, a confocal microscope (LSM 880 airy scan, Zeiss) was used in bright field with 630 × magnification. Different *MSMEG* strains were obtained during early logarithmic phase which OD_600_ around 0.5. For each strain, there were 30 random selected cells photo by Zen 2.3 (Zeiss) for length and width measurement.

For SEM, a modified method from previous studies was used (Julistiono *et al*., 2018, Fujiwara *et al*., 2015). *MSMEG* strains were mixed with 2.5% of glutaraldehyde in phosphate buffer saline (PBS) solution for sample fixation. Samples were followed by twice washing by PBS solution for 30 min, fixation by 1% OsO_4_ in PBS solution for 1 hour. Finally, dehydration progress was done with graded ethanol series (30, 50, 70, 85, 95, and 100%) for 15 min at each concentration. Cells were observed with SEM (JEOL JSM 6510) at electron energy of 30 kV. Cell length (L) and width (W) value was measured via ImageJ and cellular volume (V) was calculated with the following equation (Czerwinska-Glowka and Krukiewicz, 2021).

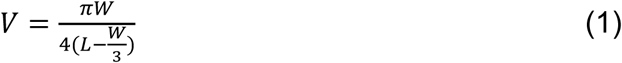

### 2.9 Colony morphology

Additional 100 μg/mL Congo red was added in 7H10 plates supplemented with 10% ADS (v/v), appropriate antibiotics to improve visualization of colony morphology (Klepp *et al*., 2012). After 72 hours, all colonies were recorded by ChemiDoc™ MP Imaging System (Bio-Rad).

### 2.10 Biofilm formation and quantification

The method for biofilm formation and quantification was adapted from previous studies with some modifications (Petchiappan *et al*., 2020, Bharti *et al*., 2020). Typically, strains were cultured for 18 hours and OD_600_ was determined, then washed twice by Sauton’s medium (REBIO) and bacteria were resuspended to an OD_600_ of 0.03 in Sauton’s medium supplemented with 2% filtered-glucose. For biofilm formation, diluted strains were cultured in polystyrene 24-well plate (Corning). All plates were sealed by plastic bags and incubated without shaking at 37 °C for 7 days until visible biofilm emerged, and imaged. For quantification, crystal violet assay was performed. Diluted strains were cultured into polystyrene 96-well plates at 37 °C for 5 days. After 5 days of culture, bacteria cultures were discarded and washed three times with sterile-water. The adherent biofilms attached to the wells were stained with 250 μl of 1% crystal violet (w/v) solution and incubated at room temperature for 15 min. The surplus dye was washed with sterile-water 4-5 times and the plate was air dried. The bound dye was finally solubilized in 250 μL 95% ethanol (v/v) and the value of OD_595_ was measured using CLARIOstar Plus (BMG Labtech).

### 2.11 Aggregation assay

The method used to study aggregation was a modified protocol from previous studies (Deshayes *et al*., 2005, Liu *et al*., 2021). Saturated cultures (OD_600_ > 2) of the strains were grown without Tween-80. Cells were resuspended in 5 mL PBS to an OD_600_ was around 1.0 and completely suspended by vortexing for 1 min. The upper part of the bacterial suspension was taken and its OD_600_ measured using a spectrophotometer (BioPhotometer plus, Eppendorf). Results at time points 0, 5 and 10 min were used for calculation of aggregation indexes using the following equation:

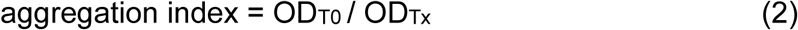

### 2.12 SDS sensitivity assays

There were two methods employed to determine SDS sensitivity. For both methods, different *MSMEG* strains were obtained during logarithmic phase and diluted to OD_600_ of 0.05 in culture medium before loading on the plates. For the first method, 7H10 agar plates supplemented with 10% ADS (v/v) were made with different concentration of SDS including 0.05%, 0.1%, 0.2% and 0.4% (w/v) according to previous study (Lefebvre *et al*., 2018) with some edition. There were 10 μL of diluted bacteria culture loaded on the plate and incubated at 37 °C for 72 hours, the sensitivity of the strains to SDS was estimated by figuring out the SDS concentration which bacteria cannot grow.

The second method was modified from a previous study (Yan *et al*., 2017), 100 μL of diluted bacterial culture was spread by a cotton swab on the 7H10 plate supplemented with 10% ADS (v/v). Sterilized Whatman discs were placed on the central of 7H10 plate, and 5 μL of different concentrations of SDS (2%, 4%, 8% and 10% (w/v)) was pipetted into the discs. All plates were incubated at 37 °C for 72 hours, the sensitivity of the strains to the SDS was estimated by measuring the diameter of the inhibition zone.

### 2.13 Minimum inhibitory concentration (MIC) assay

An MIC assay was modified from previous studies (Lu *et al*., 2003, Agrawal *et al*., 2015, Lefebvre *et al*., 2018). Strains were obtained during logarithmic phase and diluted to OD_600_ of 0.05 in culture medium before loading on the plates. Different antibiotics were diluted and mixed with 7H10 supplement with ADS during making plates. There were 10 μL of diluted bacteria culture loaded on the plate, and plates were incubated at 37 °C for 72 hours. The lowest concentration of antibiotic at which no colonies were grown was defined as MIC for the bacteria strain.

### 2.14 3D protein structure prediction and sequence alignment

The 3D protein structure prediction for full-length MSMEG_1353 was performed using SWISS-MODEL (https://swissmodel.expasy.org/) (Waterhouse *et al.,* 2018) and the AlphaFold Protein Structure Database (https://alphafold.ebi.ac.uk/) (Jumper *et al*., 2021, Varadi *et al*., 2022), employing the complete amino acid sequence of full-length MSMEG_1353. The amino acid sequences of full-length MSMEG_1353 from *MSMEG* mc^2^155 strain was obtained from the NCBI database, and this sequence was used to run protein-protein BLAST (https://blast.ncbi.nlm.nih.gov/Blast.cgi) with common bacteria and human, amino acid sequences identities higher than 25% to full-length MSMEG_1353 were selected for sequence alignment. The alignment of sequences with predicted secondary structure was obtained from ESPript (Robert and Gouet, 2014), based on 5yjz PDB model derived from the SWISS-MODEL 3D-structure prediction. Following protein homology modeling, PYMOL v. 2.3 was employed for visualizing the predicted structure of full-length MSMEG_1353. The UNIPROT (https://www.uniprot.org/) (UniProt-Consortium, 2019) was used to predict protein function or structure domain sites.

### 2.15 Sample preparation for MS

Proteome samples of the WT, aTc-induced 1353i and 1353up strains were prepared following an established method (Le *et al*., 2020) with some modifications. Cells were obtained during logarithmic phase, washed pellets were resuspended in 1 mL of 8 M urea, 1× protease inhibitor (Thermo Fisher Scientific) and 1 mM EDTA and ultrasonicated by sonication (Qsonica Q700 Sonicator with microtip probe) on an ice bath at 50% amplitude, with a pulse-on time of 5 s and a pulse-off time of 10 s for a total duration of 3 min. The lysed sample was centrifuged at 12000 × *g* for 10 min at 4 °C and supernatant was transformed into a fresh tube with extra dithiothreitol which final concentration was 5 mM. After 30 min incubation at 56 °C, extra iodoacetamide was added which final concentration was 11 mM and incubated in darkness at room temperature for another 15 min. Then, sample was mixed with 100 mM ammonium bicarbonate to dilute urea to 1 M, and protein concentration was measured for trypsin treatment via bicinchoninic acid assay (Beyotime) following its manufacturer’s recommendations.

Measured proteome sample was digested by trypsin (1 µg/µL) at a ratio of 1:50 (w/w) at 37 °C for 16 hours and trifluoroacetic acid (TFA) was added before desalting which final concentration was 0.1% (v/v). The MonoSpin C18 column (GL Sciences, Inc.) was used for proteome desalting, the centrifugation speed for whole desalting progress was 2300 × g. The column was activated with 200 µL acetonitrile (ACN) and spun for 1 min, and 200 µL of 0.1% TFA and spun for 1 min. Sample was loaded and spun for 2 min, followed with 2 min spun by 200 µL of 0.1% TFA for washing and 2 min spun by 200 µL of 60% ACN for elution. The eluted sample was dried with Savant SPD131DDA SpeedVac Concentrator (Thermo Fisher Scientific) without heating and dried sample was resuspended with 0.1% (v/v) formic acid. Samples were measured via NanoDrop 2000 spectrophotometer (Thermo Fisher Scientific) and adjusted to 0.1 mg/mL with 0.1% (v/v) formic acid for MS analysis.

### 2.16 Mass spectrometry

Six prepared peptide samples of each strain were analyzed by LC-MS according to previous studies with some modifications (Glass *et al*., 2017, Le *et al*., 2020). LC-MS was done via using an EASY-nLC 1000 (Thermo Fisher Scientific) coupled online to an LTQ-Orbitrap Elite (Thermo Fisher Scientific). Resuspended peptides were transformed into a 1.4 mL Micro-V Vial Target DP CLR (Thermo Fisher Scientific) glass sample tube and sealed by Avcs Blue cap with T/RR Septa (Thermo Fisher Scientific) seal cap. There were 10 µL suspended peptides loaded onto an Acclaim PepMap 100 C-18 precolumn (75 μm × 2 cm, particle size was 3 μm, pore size was 100 Å; Thermo Fisher Scientific), and then the precolumn was switched online with analytical Acclaim PepMap RSLC C-18 column (50 μm × 15 cm, particle size was 2 μm, pore size was 100 Å; Thermo Fisher Scientific) equilibrated in 95% solvent A (0.1% Formic Acid (v/v) in water, LC-MS Grade; Thermo Fisher Scientific) and 5% solvent B (0.1% Formic Acid (v/v) in acetonitrile, LC-MS Grade; Thermo Fisher Scientific). The peptides were eluted using solvent B at a 0.3 μl/min flow rate with the following gradient: 0 to 5% solvent B in 2 min, 5 to 30% in 150 min, 30 to 90% in 20 min and finished with 90 to 5% in 15 min.

The 10 most intense ions per survey scan were selected for higher-energy collisional dissociation fragmentation. MS was operated in the data-dependent acquisition mode with XCalibur software. Survey MS scans were acquired in Orbitrap in the 300–2000 m/z range with a resolution of 60000. The normalized collision energy for collision energy was set to 35%, the isolation width was 2.0 m/z, and the activation time was 10 ms. Three data groups of bottom-up proteomics have been deposited to the ProteomeXchange Consortium via the PRIDE partner repository (Perez-Riverol *et al*., 2022) with the dataset identifier PXD048724, PXD048725 and PXD048860, respectively.

### 2.17 Protein identification and quantification

The raw data files generated by MS were processed as described (Ganief *et al*., 2018, Le *et al*., 2020) with some modifications. The raw MS files were analyzed using MaxQuant version 1.6.7.0, and the resulting data were searched against the *MSMEG* entries of the SWISSPROT protein database [strain ATCC 700084/mc2155 UNIPROT IDs UP000000757 with a total of 6,602 entries] employing the Andromeda search engine. MaxQuant was employed with a fragment ion mass tolerance of 20 ppm for the purpose of search. In MaxQuant, carbamidomethyl of cysteine was designated as a fixed modification, while oxidation of methionine, acetyl of the N-terminus, and phosphorylation of serine and threonine were specified as variable modifications.

To validate MS/MS based peptide and protein identifications, Scaffold (version Scaffold_5.0.0, Proteome Software Inc.) software was utilized. Peptide identifications were considered acceptable if they could be confidently established at a probability greater than 6.0% to achieve a false discovery rate less than 1.0%, employing the Peptide Prophet algorithm (Keller *et al*., 2002) with Scaffold delta-mass correction for accurate results assessment. Protein identifications were considered valid if they could be confidently established with a probability exceeding 61.0% to achieve a false discovery rate of less than 1.0%, and if they were supported by the identification of at least two peptides. The assignment of protein probabilities was performed using the Protein Prophet algorithm (Nesvizhskii *et al*., 2003). Proteins containing similar peptides that cannot be distinguished solely based on MS/MS analysis were grouped together in accordance with the principles of parsimony. Proteins sharing significant peptide evidence were clustered accordingly. The protein amount of the WT strain was utilized as a standard for data normalization and proteomes of the 1353up and aTc-induced 1353i strains were compared with normalized WT proteome to figure out protein expression level shift. Prior to analysis, the normalized data from each strain within the same experiment were averaged (Cho *et al*., 2012, Wolfe *et al*., 2013).

### 2.18 Statistical analysis

All experiments were performed at least three independent replicates, analyzed and presented as mean ± SD via GraphPad Prism v8.0.1. A two-tailed unpaired *t*-test was used for two groups to detect Statistically significant. Significantly different was determined when *p* value was less than 0.05.

## 3. Results

### 3.1 *MSMEG_1353* expression levels in engineered strains

First, strains in which the essential *MSMEG_1353* gene is overexpressed or could be knocked down were constructed. *MSMEG_1353* was amplified from the mc^2^155 genome by PCR and ligated into the pMV261 plasmid harboring the constitutively active promoter *Mycobacterium bovis hsp60*. Empty pMV261 and pMV261-1353 were transformed into competent *MSMEG* cells to generate a negative control strain (designated WTP) and an *MSMEG_1353* overexpression strain (designated 1353up), respectively. Both strains and the WT harboring no plasmid were grown in liquid 7H9 medium for up to 72 hours and subjected to RT-qPCR to examine their mRNA expression level of *MSMEG_1353* (Fig. 1A). As expected, the 1353up strain expressed *MSMEG_1353* at 100-fold (*p* < 0.01) higher levels compared to both control strains in samples cultured for 18 hours, and at 56-fold (*p* < 0.05) higher levels after 72 hours. These results indicate that the 1353up strain can be utilized for further characterization of the role of *MSMEG_1353* overexpression for at least up to 72 hours after the start of a culture.

**Fig. 1.**
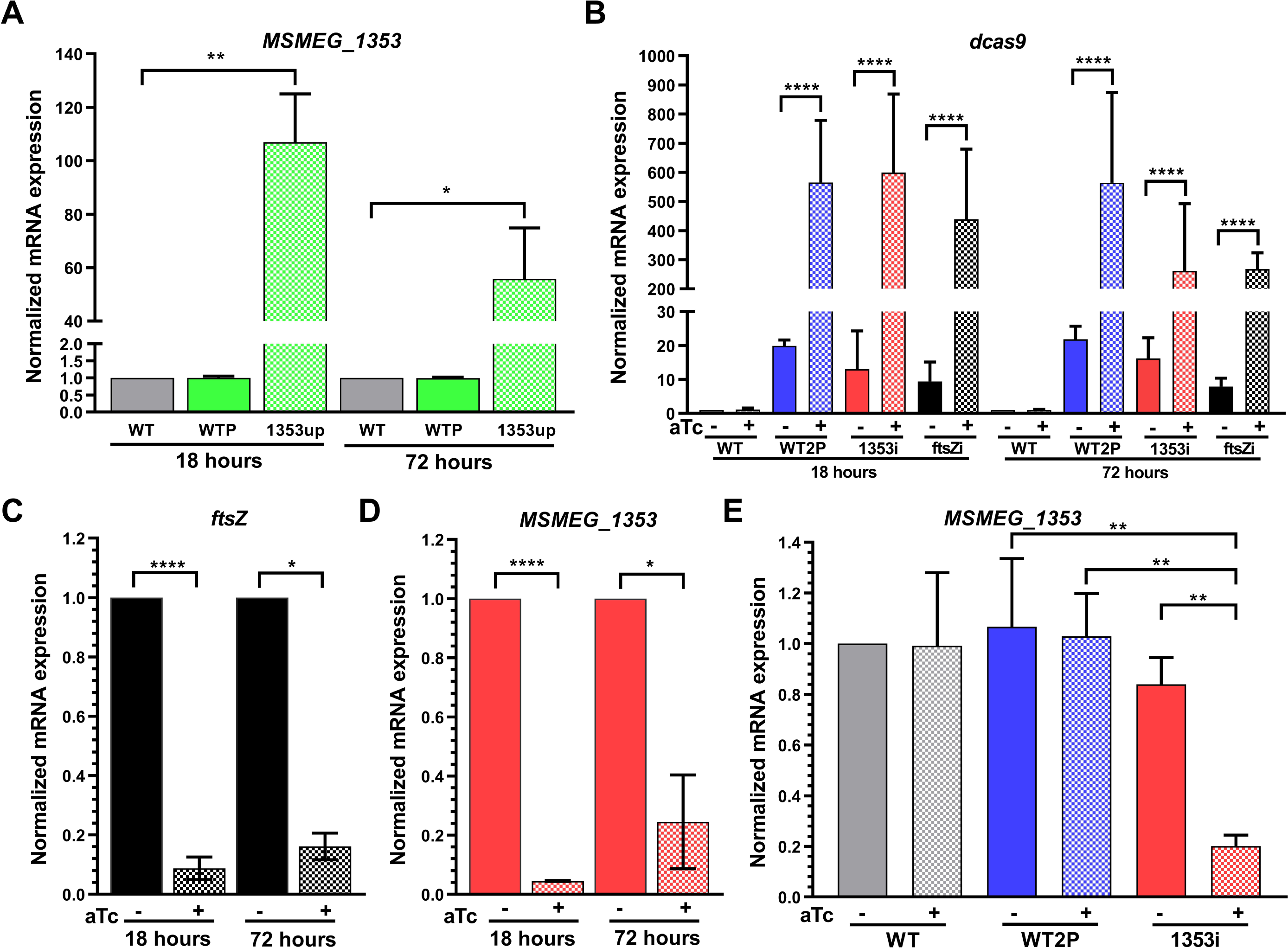
Expression of target genes in mutant strains. Relative mRNA expression was determined using the 2^−ΔΔCT^ method with *rpoD* as the reference gene. **(A)** *MSMEG_1353* expression in the WTP and 1353up strains normalized to the WT strain. **(B)** Expression of *dcas9* normalized to the WT strain in the presence or absence of aTc. Expression of **(C)** *ftsZ* in the ftsZi strain and **(D)** *MSMEG_1353* in the 1353i strain after 18 and 72 hours in the presence or absence of aTc. **(E)** *MSMEG_1353* expression normalized to the WT strain after 18 hours in the presence or absence of aTc. Results are the mean of three independent experiments performed in triplicate each, and the error bars represent ± standard deviation. Asterisks indicate significant differences in an unpaired t test (* *p* < 0.05; ** *p* < 0.01; *** *p* < 0.001).

Knockdown strains that make use of an inducible dead *cas9* gene (*dcas9*) in a CRISPRi system were constructed by introducing two plasmids into *MSMEG* (Singh *et al*., 2016). The first is pRH2502, which contains *dcas9* under the control of the TetR-regulated uvtetO promoter which is induced by addition of aTc. The second plasmid transformed into *MSMEG* was pRH2502, with either *MSMEG_1353* or *ftsZ* (as a CRISPRi system positive control) cloned behind the TetR-regulated smyc. This resulted in strains that were designated as 1353i and ftsZi, respectively. To exclude any influence of the plasmids used for CRISPRi, a negative control strain, named WT2P, was made by introduction of pRH2502 and an empty pRH2521. These strains were cultured in the presence or absence of aTc for up to 72 hours and subsequently subjected to RT-qPCR to examine the expression levels of *dcas9*, *MSMEG_1353* and *ftsZ*.

Without aTc induction, low levels of *dcas9* mRNA were observed in the WT2P, 1353i, and ftsZi strains (Fig. 1B), with varying levels of expression between 0.1 and 0.3 relative to *rpoD*. This transcription in these uninduced CRISPRi strains will be referred to as “leaky expression”. In the aTc-induced strains, a 10- to 69-fold (*p* < 0.0001) increase in *dcas9* mRNA expression level was measured at both 18- and 72-hours compared to the non-induced CRISPRi strains. The mRNA expression of *ftsZ* in the aTc-induced ftsZi strain (Fig. 1C) and of *MSMEG_1353* in the aTc-induced 1353i strain (Fig. 1D) were normalized to the uninduced strains. After 18- and 72-hours culture, the expression level of *ftsZ* was suppressed to 0.09 and 0.16, respectively. Similarly, the expression level of *MSMEG_1353* was inhibited in the aTc-induced 1353i strain to 0.04 and 0.25 of the levels in the uninduced 1353i strain, respectively. These results indicate that the aTc-induced 1353i strain can be utilized for characterization of the role of *MSMEG_1353* for at least up to 72 hours after the start of a culture. The leaky expression of *dcas9* observed in the uninduced 1353i strain resulted in about 20% inhibition of *MSMEG_1353* expression compared to the WT (Fig. 1E), and thus this strain will be useful to investigate whether intermediate levels of *MSMEG_1353* display phenotypes deviating from the WT.

### 3.2 Morphology changes in the *MSMEG_1353* knockdown

First, we investigated whether changed levels of MSMEG_1353 result in any growth defects or morphology changes. Strains were grown in the presence or absence of aTc for up to 60 hours and OD_600_ was measured every three hours in a spectrophotometer (Fig. 2A). The *ftsZ*-deficient *MSMEG* did not show any increase in OD_600_ after induction with aTc (data not shown), as was expected because cell division in bacteria lacking FtsZ will be stalled (Singh *et al*., 2016). The *MSMEG_1353* overexpression strain did not display any significant difference compared to the WT strain (data not shown). Both aTc treated and non-treated cultures of the WT2P strains exhibited a delay in log-phase initiation of around 1 hour compared to the WT strain. This result suggests that the presence of CRISPRi plasmids may exert a subtle influence on the growth of *MSMEG*. Both the 1353i and aTc-induced 1353i strains exhibited an even more pronounced delayed entry into log-phase compared to the aTc-induced WT2P strain, pointing to a role for the lack of *MSMEG_1353*. The uninduced 1353i strain showed a delay of 3 hours, while this further increased to 5-7 hours in independent experiments with the aTc-induced 1353i strain. Apparently, the lack of *MSMEG_1353* considerably postpones the initiation of log-phase, with this delay being positively correlated to the degree of inhibition of *MSMEG_1353* expression. After initiation of the exponential growth phase, the doubling time of lower levels of MSMEG_1353 was 1.2-fold (*p* < 0.05) when comparing with the WT strain, while it was similar as the value from its negative control strains and the final OD_600_ were comparable.

**Fig. 2.**
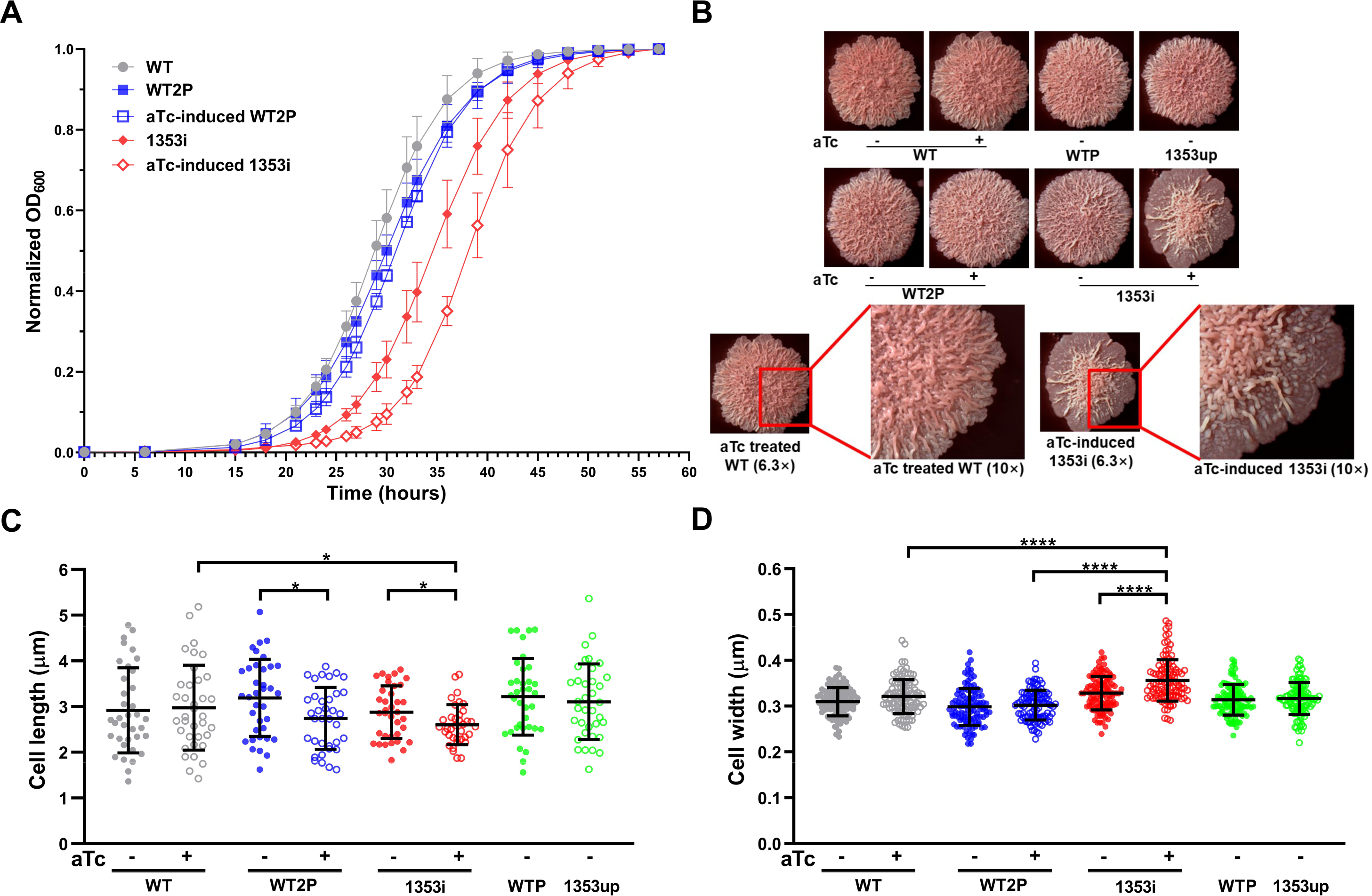
Growth-related characteristics and cellular dimensions of strains with differential *MSMEG_1353* expression. **(A)** Growth curve of strains grown in the presence or absence of aTc. **(B)** Colony morphology on 7H10 plates supplemented with Congo red after 72 hours of culture in the presence or absence of aTc. **(C)** Length and **(D)** width of 36 individual bacteria from strains grown in the presence or absence of aTc. Cells were randomly taken from multiple SEM images. Results are the mean of three independent biological experiments performed in triplicate each, and the error bars represent ± standard deviation. Asterisks indicate significant differences in an unpaired t test (* *p* < 0.05; **** *p* < 0.0001).

Next, we investigated the impact of MSMEG_1353 on the cell envelope by utilizing Congo red, as this enhances visualization of colony morphology and facilitates evaluation of changes in lipid and lipoprotein contents within the cell envelope (Cangelosi *et al*., 1999, Lefebvre *et al*., 2018). All colonies exhibited similar sizes on the 7H10 plate supplement with Congo red after a 3-day incubation period, irrespective of MSMEG_1353 levels. The color picture of colony samples of the eight strains were observed (Fig. 2B), with no discernible color variations visible to the naked eye, suggesting that MSMEG_1353 did not influence the hydrophobicity of *MSMEG* in these experiments. The WT strain colony showed a uniform density of corrugation across the surface of the colony, whereas strains treated with aTc display wider and more jagged edges compared to the non-treated strains. Interestingly, on the colony surface of the aTc-induced 1353i strain, a remarkable difference could be observed, with dense corrugation in the central region and almost no corrugation observed at the colony surface edge.

The eight strains were grown in liquid 7H9 medium until early log-phase and subsequently subjected to phase confocal microscopy (Fig. S2) and SEM for morphological inspection (data not shown) and measurement of cellular dimensions (Fig. 2C, 2D). Downregulating *ftsZ* gene expression levels caused a 4.2-fold (*p* < 0.05) elongation in the aTc-induced ftsZi strain compared to the WT strain by phase contrast microscopy, showing very long cells with multiple branches which aligns with the requirement of FtsZ for cell division (Kang *et al*., 2008), and indicating that the CRISPRi system is functional. Interestingly, significant changes in cellular dimensions were observed. An increase in width of 19% for the aTc-induced 1353i strain was apparent. WT cells were 2.92 ± 0.93 μm in length and 0.31 ± 0.03 μm in width, in line with literature for *MSMEG* (Zou *et al*., 2017). Unexpectedly, all aTc-induced CRISPRi strains displayed a slight decrease in length when compared to their uninduced counterpart strains. Cells from the aTc-induced 1353i strain were the shortest of all (Fig. 2C), but the mean value (2.6 ± 0.44 µm) was not significantly different from the aTc-induced WT2P strain (2.7 ± 0.67 µm), which prevents us to conclude that the reduction in length can solely be attributed to a deficiency of MSMEG_1353. In line with phase contrast microscopy results (Fig. S2), decreased levels of MSMEG_1353 in the aTc-induced 1353i strain consistently led to an increase in cell width of around 10% (Fig. 2D).

The length-to-width ratio (Fig. S3) and cellular volume (Fig. S4) for each strain were calculated. All aTc-treated strains show a lower ratio value, especially in the CRISPRi strains, suggesting that aTc treatment and CRISPRi plasmids activation may influence this change. This should be considered in future work. Interestingly, the aTc-induced 1353i strain has the lowest ratio value (between 0.8 and 0.9-fold (*p* < 0.05)) and the largest cellular volume (around 1.3-fold (*p* < 0.001)) compared to the control strains suggesting that lower levels of *MSMEG_1353* influences cell shape, likely via changes in its cell envelope.

### 3.3 Reduction in biofilm formation and aggregation in the *MSMEG_1353* knockdown

Changes in the lipid composition of the cell wall can exert a substantial impact on surface hydrophobicity and diverse cellular surface properties, such as biofilm/pellicle formation and cellular aggregation (Singh. *et al*., 2016). The pellicles are hypothesized to be biofilm-like structures that assemble at the air-liquid interface. The presence of thick and robust pellicles is indicative of a higher bacterial biofilm formation ability (Singh. *et al*., 2016). When cultured in polystyrene plates, it was observed that the aTc-induced 1353i strain was hardly able to form a biofilm on the air-liquid interface in 24-wells plates after 7 days, while other strains formed a clear biofilm with some corrugations in the pellicles within 5 days (Fig. 3A). Biofilm formation was further quantified in a polystyrene 96-well plate using 1% crystal violet (Fig. 3B). Crystal violet is binding to the hydrophobic moieties in the mycobacterial cell envelope which can be quantified using absorbance at OD_595_. The WT strain showed an average OD_595_ value of 0.65. The aTc-induced 1353i strain was binding much less crystal violet, indicated by 3.3-fold lower value (*p* < 0.001) compared to its negative controls, and the uninduced 1353i strain showed values that were 20% lower than the WT due to its leaky expression. These results suggest that MSMEG_1353 is involved in production of hydrophobic compounds needed for biofilm formation.

**Fig. 3.**
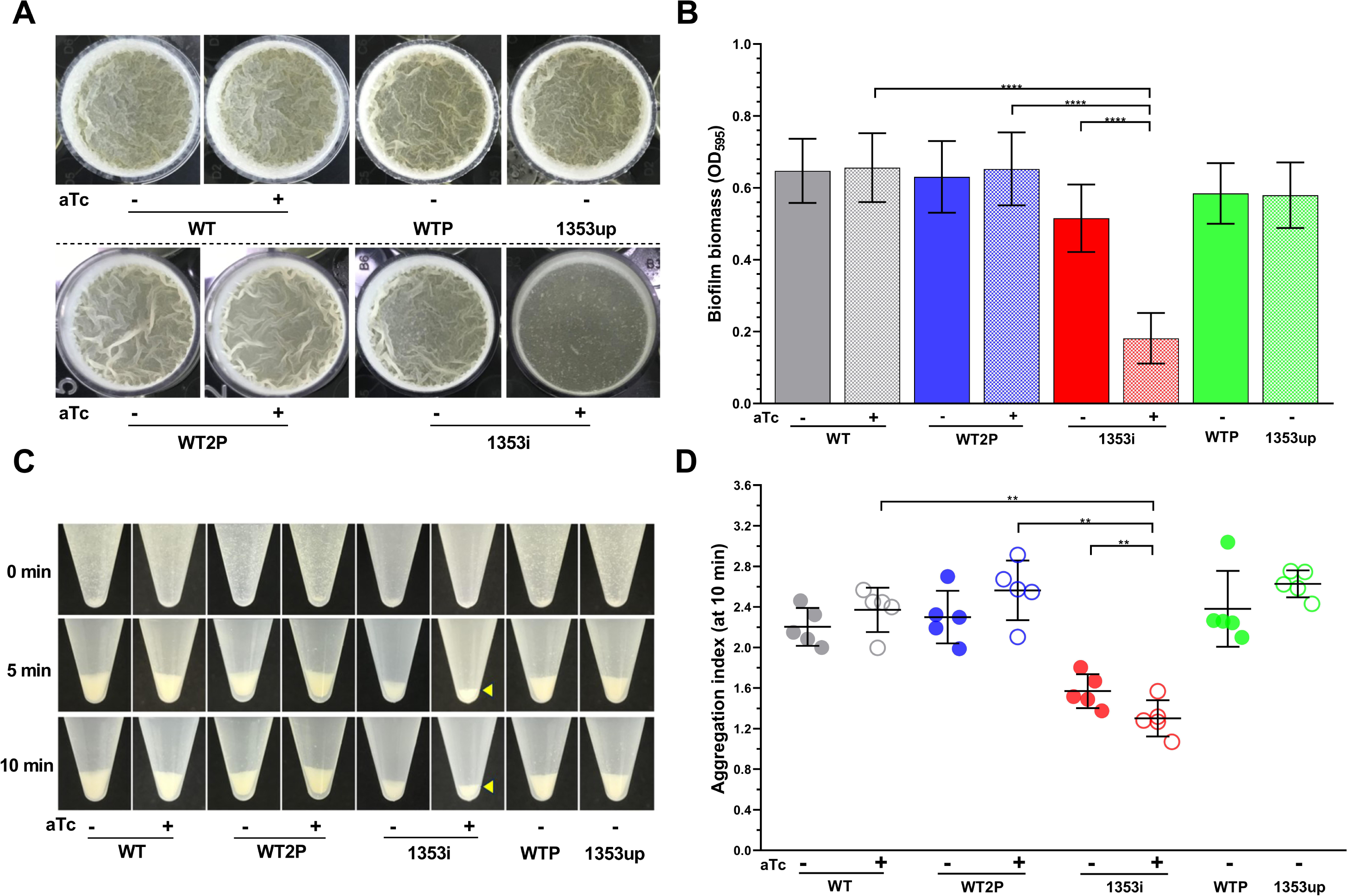
Biofilm formation and aggregation ability of strains with differential *MSMEG_1353* expression. **(A)** Top view of pellicle formation at the air-liquid interface by strains grown in polystyrene 24-well plates in Sauton’s medium in the presence or absence of aTc for six days. **(B)** Quantification of biofilm formation by crystal violet staining of strains grown in polystyrene 24-well plates in Sauton’s medium in the presence or absence of aTc for five days. **(C)** Aggregation by strains grown in PBS in the presence or absence of aTc in 15 mL polyethylene tubes. Yellow arrows indicate the smallest pellet in the aTc-induced 1353i strain at different time points of settling after vigorous vortexing. **(D)** Aggregation index at t = 10 min of strains grown in 7H9 medium without Tween-80 for 2 days in the presence or absence of aTc. Results are the mean of five independent biological experiments performed in triplicate each, and the error bars represent ± standard deviation. Asterisks indicate significant differences in an unpaired t test (* *p* < 0.05; ** *p* < 0.01).

Under standard laboratory culture conditions without detergent, *MSMEG* can spontaneously aggregate and form hydrophobic clumps (Sani *et al*., 2010, Julian *et al*., 2010). Aggregation is believed to precede maturation of biofilms and formation of pellicles in *Mycobacteria* (Bharti *et al*., 2020). Aggregation speed among all eight strains was compared at 0, 5, and 10 min (Fig. 3C) by observation with the naked eye. The WT and other strains show a clear aggregated pellet and no observed changes after 5 min, while both the uninduced 1353i and aTc-induced 1353i strains exhibited fewer aggregated cells even after an incubation period of 18 hours compared to all other strains. Additionally, the aTc-induced 1353i strain displayed a lower number of aggregated cells compared to the uninduced 1353i strain. Both the uninduced 1353i and aTc-induced 1353i strains also had a cloudier upper aqueous layer than the other strains. However, relying solely on visual inspection may not provide accurate results. Hence, the aggregation index was utilized, a lower aggregation index indicating less cellular aggregation. The results of the aggregation index indicate that both the uninduced 1353i and aTc-induced 1353i strains exhibited reduced cellular aggregation compared to other strains after 5 min (Fig. S5) and 10 min (Fig. 3D). The aTc-induced 1353i strain showed an aggregation index of 0.55-fold (*p* < 0.01) compared to the aTc-induced WT2P strain and 0.85-fold (*p* < 0.01) compared to the uninduced 1353i strain. Clearly, the lower level of MSMEG_1353 leads to a lower rate of cell aggregation.

### 3.4 Increased sensitivity to SDS and antibiotics when MSMEG_1353 is downregulated

SDS can be utilized to test the integrity of the bacterial cell wall (Meng *et al*., 2017), simulating surfactant stress encountered by *Mtb* during infection (Yan *et al*., 2017). A disc diffusion assay with different concentration of SDS was performed (Fig. S6). A 1.3 to 1.7-fold (*p* < 0.01) increase in diameter of the inhibition zone was observed for the aTc-induced 1353i compared to the control strains across all SDS concentrations tested (Fig. 4A). The increased sensitivity of SDS when MSMEG_1353 is lacking was also apparent when the strains were grown on 7H10 plates supplemented with different concentrations of SDS, with the aTc-induced 1353i strain failing to form colonies on a plate containing 0.02% SDS in 72 hours (Fig. S7).

**Fig. 4.**
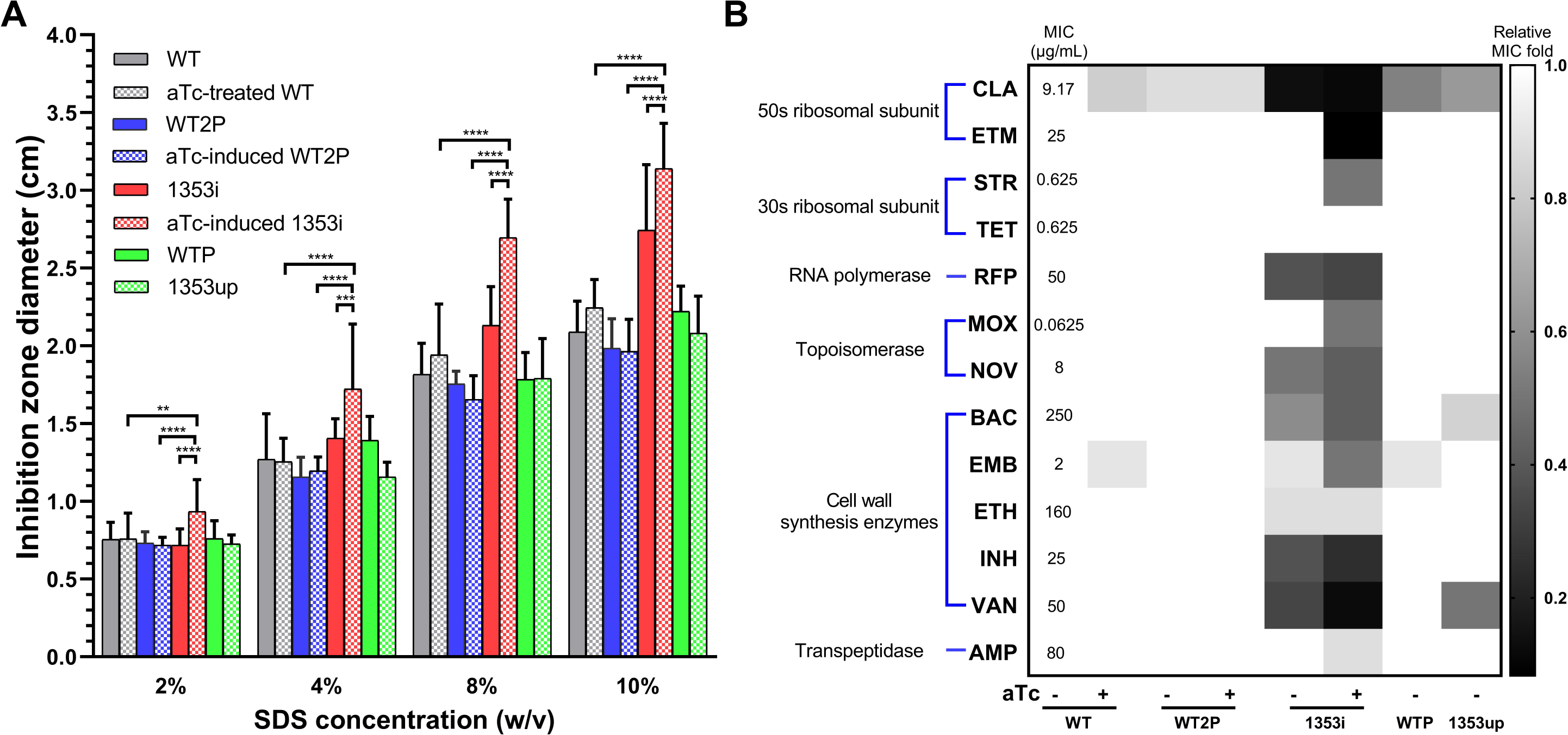
Inhibition of strains with differential *MSMEG_1353* expression by SDS and antibiotics. **(A)** Diameter of inhibition zones of strains grown in the presence or absence of aTc on 7H10 plates for discs containing different concentrations of SDS. **(B)** The MICs for 13 antibiotics of the untreated WT strain are shown in numerical values and were used for normalization (CLA-clarithromycin; ETM-erythromycin; STR-streptomycin; TET-tetracycline; RFP-rifampicin; MOX-moxifloxacin; NOV-novobiocin; BAC-bacitracin; EMB-ethambutol; ETH-ethionamide; INH-isoniazid; VAN-vancomycin; AMP-ampicillin). The grey scale on the right indicates the relative fold change in MIC for the other strains grown in the presence or absence of aTc. Antibiotics are combined into 6 groups based on their molecular target. Results are the mean of three independent biological experiments performed in triplicate each, and the error bars represent ± standard deviation. Asterisks indicate significant differences in an unpaired t test (** *p* < 0.01; *** *p* < 0.001; **** *p* < 0.0001).

Next, the MIC of several antibiotics were employed to interrogate the integrity of the cell envelope by culturing our strains on 7H10 plates with various concentrations of several. *MSMEG* has intrinsic resistance to macrolides and isoniazid (Nash, 2003, Teng and Dick, 2003), and is susceptible to tetracycline antibiotics and fluoroquinolones (Crowley *et al*., 2024). A total of thirteen antibiotics were tested in MIC assays (Fig. 4B), and the observed MIC values of the WT strain fall within the same range as previous studies (Lu *et al*., 2003, Baptista *et al*., 2018, Rai and Mehra, 2021). Ten of the thirteen antibiotics antibiotics displayed a 50% or more reduction in MIC values on the aTc-induced 1353i strain compared to the WT strain. The three antibiotics for which the sensitivity was hardly or not affected were tetracycline, ampicillin and ethionamide; all antibiotics with a relatively small molecular weight. The downregulated MSMEG_1353 renders *MSMEG* more vulnerable to the larger antibiotics, indicating a reduced selective permeability of the cell envelope, enabling these antibiotics to reach their targets more easily.

### 3.5 Sequence alignment and computational 3D structure prediction showed potential kinase-like active site residues

There were nine amino acid sequences identified via protein-protein BLAST that matched in a range of identities higher than 25% to full-length MSMEG_1353 (Table S2), all of them containing a predicted protein kinase-like domain according to UNIPROT. Amino acid sequence alignment showed high conservation of 25 amino acids, primarily in regions 73-76, 130-167, 204-209, 291-296, and 309-311 when compared to orthologous proteins (Fig. S8). According to the UNIPROT database, the active site or substrate binding region of UbiB from *E. coli* was located at conserved positions L130-V138 and K153 as ATP binding sites and D288 as proton acceptor (UniProt-Consortium, 2019, Madeira *et al*., 2019). It was likely that the corresponding amino acids also comprise the active site in MSMEG_1353 (*i.e*., 132F-140V, 154K and 291D).

AlphaFold predicted a 3D structure for full-length MSMEG_1353 with an average pLDDT score of 77.38 (Fig. S9). The pLDDT score corresponds to the model’s prediction on the local Distance Difference Test (lDDT-Cα), with a range from 0 to 100 where higher scores indicate greater confidence (Varadi *et al*., 2024). Both the C- and N-terminus regions of this structure exhibit low pLDDT scores (< 50), indicating flexible regions. However, the central sequence of full-length MSMEG_1353 region in this structure displays high pLDDT scores (> 70), containing a majority of conserved amino acids. Thus, the conserved amino acids within residues 130-167, 204-209, 291-296, and 309-311 are potentially part of the active site which is fitting the ATP binding region prediction from UNIPROT (H121-E448). Among them, S136, Q139, G148, N296, and G311 represent uncharged polar amino acids; R141 and K154 are positively charged polar amino acids; while D167, E204, E209, D291, and D309 are negatively charged and polar. These amino acids could be tested by site-directed mutagenesis to identify the active site related residues and confirm kinase activity of MSMEG_1353.

### 3.6 Influence of MSMEG_1353 expression on the proteome

Using six replicates for each MS analysis, 1754 proteins were identified in the WT strain, while 1592 and 1571 proteins were identified in the aTc-induced 1353i and 1353up strains, respectively. A volcano plot (Wolfe *et al*., 2013) displays upregulation and downregulation of all identified proteins, in either the aTc-induced 1353i or 1353up strain, compared to the WT strain (Fig. S10). The amount of MSMEG_1353 was decreased 2-fold (*p* < 0.01) in the aTc-induced 1353i strain and increased 8-fold (*p* < 0.05) in the 1353up strain compared to the WT strain, in line with what could be expected based on the observed mRNA expression changes.

Seven proteins showed the most interesting and profound changes in expression as a result of the lack of MSMEG_1353 when the complete data set was screened using the following criteria (Table 3). Firstly, they displayed more than 1.5-fold increased or decreased expression in either the aTc-induced 1353i or 1353up strain compared to the WT strain (p-value < 0.05). Furthermore, their expression profile was opposite in the aTc-induced 1353i and 1353up; implicating that the amount of MSMEG_1353 in the cell is an important factor. MSMEG_0194, MSMEG_0806, MSMEG_1049 and MSMEG_3945 were predicted as serine esterase in cutinase family protein, a peptide hydrolase, a methyltransferase involved in amino acid metabolism and a universal stress protein family protein, respectively, based on their homologous proteins according to UNIPROT. MSMEG_0710 (GrpE) is an indispensable component of the Mycobacterial cell and functions as a nucleotide exchange factor which enables ATP binding to initiate protein folding (Lupoli *et al*., 2016). MSMEG_0911 (ICL) is the predominant one of two isocitrate lyase isozymes, playing a pivotal role in growth on acetate or fatty acid as the exclusive carbon and energy source (Ko *et al*., 2021). ICL is known as the first enzyme in the glyoxylate cycle and is upregulated under hypoxic conditions in *Mtb* (Gould *et al*., 2006). MSMEG_0913 is an enzyme in mycolic acid biosynthesis that leads to the creation of trans counterparts (Laval *et al*., 2008).

**Table 3.** P roteins selected from MS experiments.

| ID in MSMEG | Enzyme name | Reference |
| --- | --- | --- |
| MSMEG_0194 | serine esterase, cutinase family protein | - |
| MSMEG_0710<br>(GrpE) | co-chaperone GrpE | (Lupoli *et al.*, 2016) |
| MSMEG_0806 | peptide hydrolase | - |
| MSMEG_0911 (ICL) | isocitrate lyase | (Ko *et al.*, 2021) |
| MSMEG_0913 | methoxy mycolic acid synthase 1 | (Laval *et al.*, 2008) |
| MSMEG_1049 | methyltransferase | (Rokicki *et al.*, 2021) |
| MSMEG_3945 | universal stress protein family protein | - |

### 3.7 Validation by RT-qPCR of increased expression of isocitrate lyase when MSMEG_1353 is low

To confirm these observed changes in protein expression levels, RT-qPCR was performed on the genes encoding these seven proteins (Fig. 5). All strains were paired into six comparison pairs for single factor impact on target gene expression and the ratio between relative expression values for these strain pairs was computed. The changes in expression levels for MSMEG_1353 served as an obvious control and were consistent with expectations. High and thus unreliable ct values were found in *MSMEG_0806*, although the result was in line with the MS data. The RT-qPCR data for *MSMEG_0194*, *MSMEG_0710*, *MSMEG_0913*, *MSMEG_1049* and *MSMEG_3945* result was either inconclusive or not significant. The only gene that exhibited a similar and significant expression profile in the RT-qPCR as the MS experiment was *MSMEG_0911* (*icl*), with an elevated ratio (4.8-fold (*p* < 0.05)) in the aTc-induced 1353i and aTc-induced WT2P strains pair. This gives us the confidence to state that the expression of *MSMEG_0911* (*icl*) is negatively correlated with the presence of MSMEG_1353, thereby suggesting a functional link between both proteins.

**Fig. 5.**
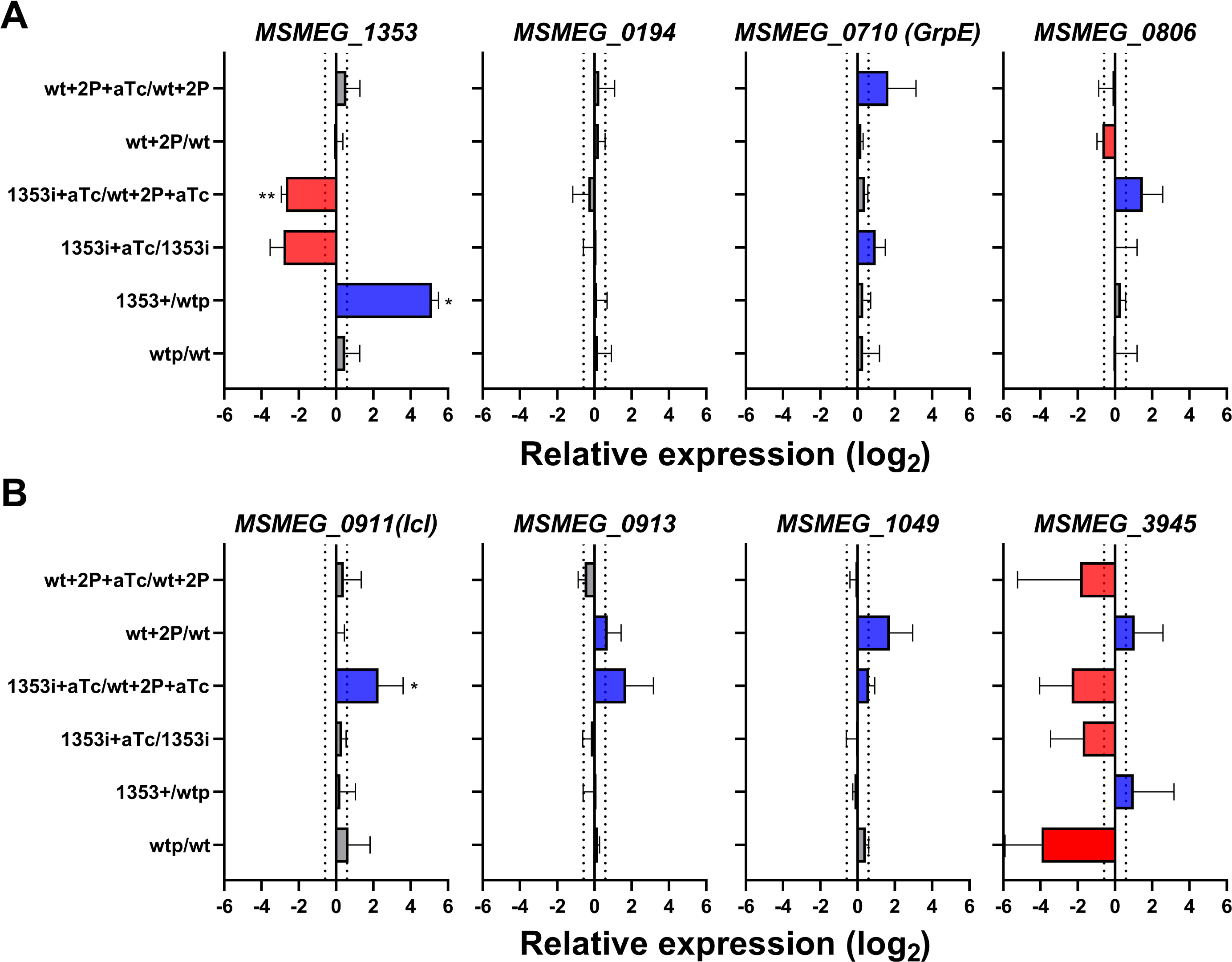
Relative expression levels of potential target genes subjected to *MSMEG_1353* regulation. The log_2_ fold change values of mRNA relative expression levels of selected genes when compared between strain pairs. Red bars indicate mRNA relative expression level was downregulated; blue bars signify mRNA relative expression level was upregulated. Grey bars represent genes that exhibit a fold change lower than 1.5. Results are the mean of three independent biological experiments performed in triplicate each, and the error bars represent ± standard deviation. Asterisks indicate significant differences in a one sample t test (* *p* < 0.05; ** *p* < 0.01).

### 3.8 *FAS* genes expression influenced by the downregulation of MSMEG_1353

Besides for validation of the MS results, we also used RT-qPCR to investigate the expression of a set of genes involved in lipid synthesis by looking at the impact of MSMEG_1353 levels on *FAS-I* and *FAS-II* genes, respectively (Fig. 6A). When the amount of *MSMEG_1353* present in the cell was low, the expression of *MSMEG_4326* (1.9-fold (*p* < 0.05)), *MSMEG_4328* (1.5-fold (*p* < 0.05)) and *MSMEG_4329* (2.1-fold (*p* < 0.05)) was upregulated, and *MSMEG_4325* and *MSMEG_4327* were more than 2.0-fold upregulated (albeit with a high standard deviation). Notably, similar trends in changes to protein amounts fold change in MS results (Table S3) were observed for alterations in gene expression levels of *MSMEG_4325-4328* (note, MSMEG_4329 was not detected) under both up- and down-regulated conditions of *MSMEG_1353*, suggesting a directly or indirectly negative correlation between *MSMEG_4325-4329* expressions regulated by this particular gene.

**Fig. 6.**
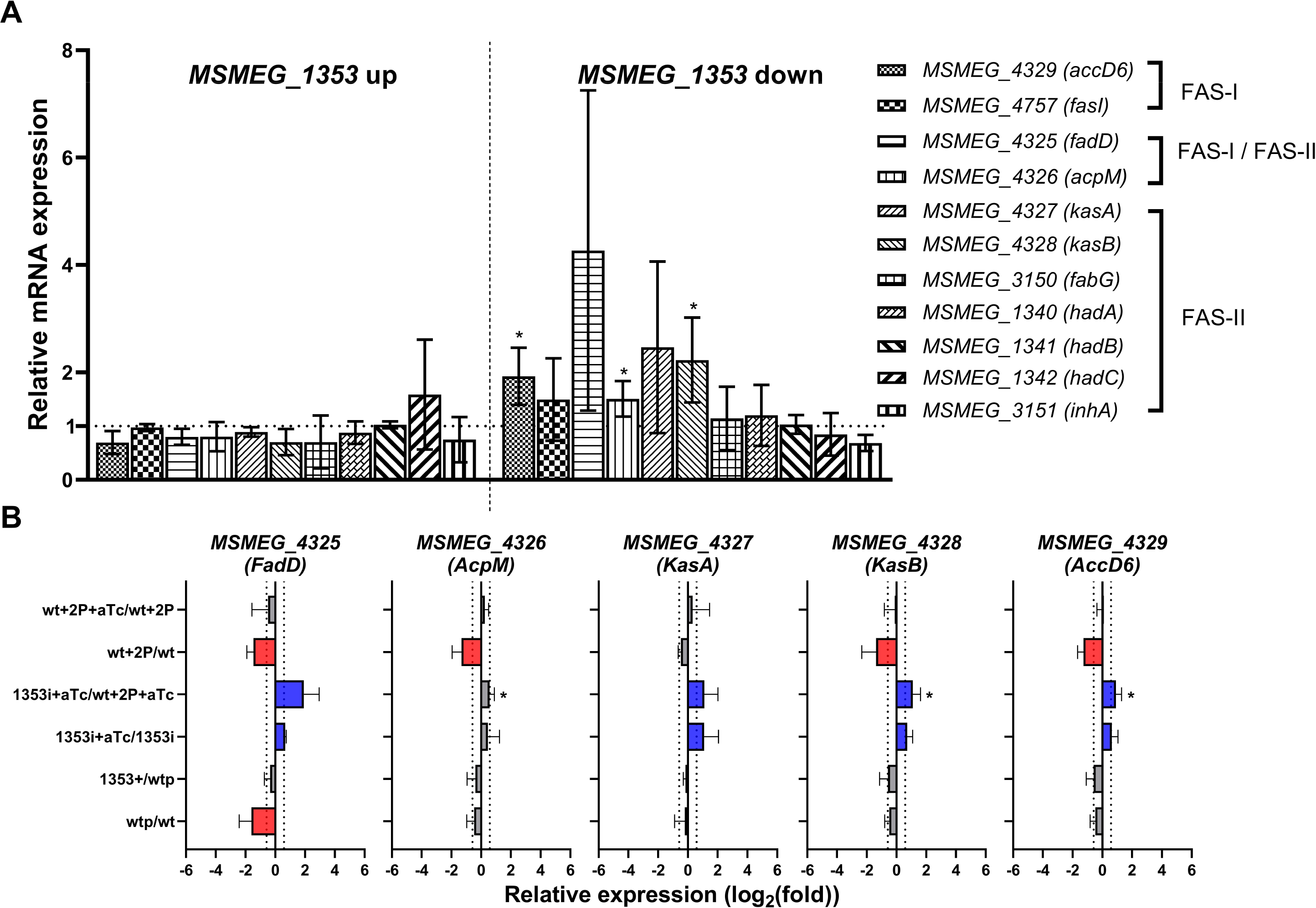
Normalized relative expression of FAS genes subjected to *MSMEG_1353* regulation and relative expression folds of selected genes between strain pairs. **(A)** The horizontal dotted line indicates no effect by *MSMEG_1353* up- or down-regulation on expression of FAS-I and FAS-II related genes. **(B)** The log_2_ fold change values of relative mRNA expression levels of *FAS* genes compared between strain pairs. Red bars indicate mRNA relative expression level was downregulated; blue bars signify mRNA relative expression level was upregulated. Grey bars exhibit a fold change lower than 1.5. Results are the mean of three independent biological experiments performed in triplicate each, and the error bars represent ± standard deviation. Asterisks indicate significant differences in a one sample t test (* *p* < 0.05; ** *p* < 0.01).

Expression ratios of *MSMEG_4325-4329* between control strains are shown in Figure 6B, clearly demonstrating the inverse correlation with MSMEG_1353 expression levels. Interestingly, a 2.1-fold (*p* = 0.17) overexpression of *MSMEG_4327* was observed in the *MSMEG_1353* downregulated strain. *MSMEG_4328-4329* showed an approximately 2.0-fold (*p* < 0.05) overexpression and *MSMEG_4326* showed a 1.4-fold (*p* < 0.05) overexpression when *MSMEG_1353* was downregulated, indicating that *MSMEG_4325-4329* were upregulated when MSMEG_1353 was lowered in the cell. These data suggest MSMEG_1353 may be a suppressor of this operon or it has an indirect effect via a suppressor that was removed due to the downregulation of MSMEG_1353. A role for MSMEG_1353 in lipid synthesis seems very likely based on its effect on all genes assessed by RT-qPCR.

## 4. Discussion

All data obtained in this study point to a role for MSMEG_1353 in regulation of genes that are involved in stress response and/or cell envelope integrity, in particular lipid synthesis. We have outlined the possible interplay between MSMEG_1353 and several genes that were identified during this work (Fig. 7). In this section we will discuss the proposed model in the light of the experimental data. Both the aTc-induced and uninduced 1353i strains exhibit a delayed initiation of log phase growth of approximately 5-7 hours and around 3 hours compared to the aTc-induced WT2P strain, respectively. The delay in growth was positively correlated with the expression levels of MSMEG_1353, and no changed growth patterns were observed in any of the control strains. It has been shown previously that lower levels of mycolic acid biosynthesis proteins within *Mycobacteria* can consistently lead to delays in log-phase starting time ranging from hours to 2 days, notably for HadD (a potential hydratase/dehydratase in FAS-II) (Lefebvre *et al*., 2018), methyltransferases for mycolic acid cyclopropanation (CmaA2 as example) (Barkan *et al*., 2009), an arylamine *N*-acetyltransferase (Bhakta *et al*., 2004), and a short-chain dehydrogenase (MSMEG_4722) (Bhatt *et al*., 2008). In our SEM analysis, an increase of approximately 10% in cell width and 1.3-fold in cellular volume were observed in the aTc-induced 1353i strain compared to the WT strain. Such a phenotype has been observed previously as the result of an increase in mycolic acid carbon-chain length (Chen *et al*., 2020, Zhang *et al*., 2022).

**Fig. 7.**
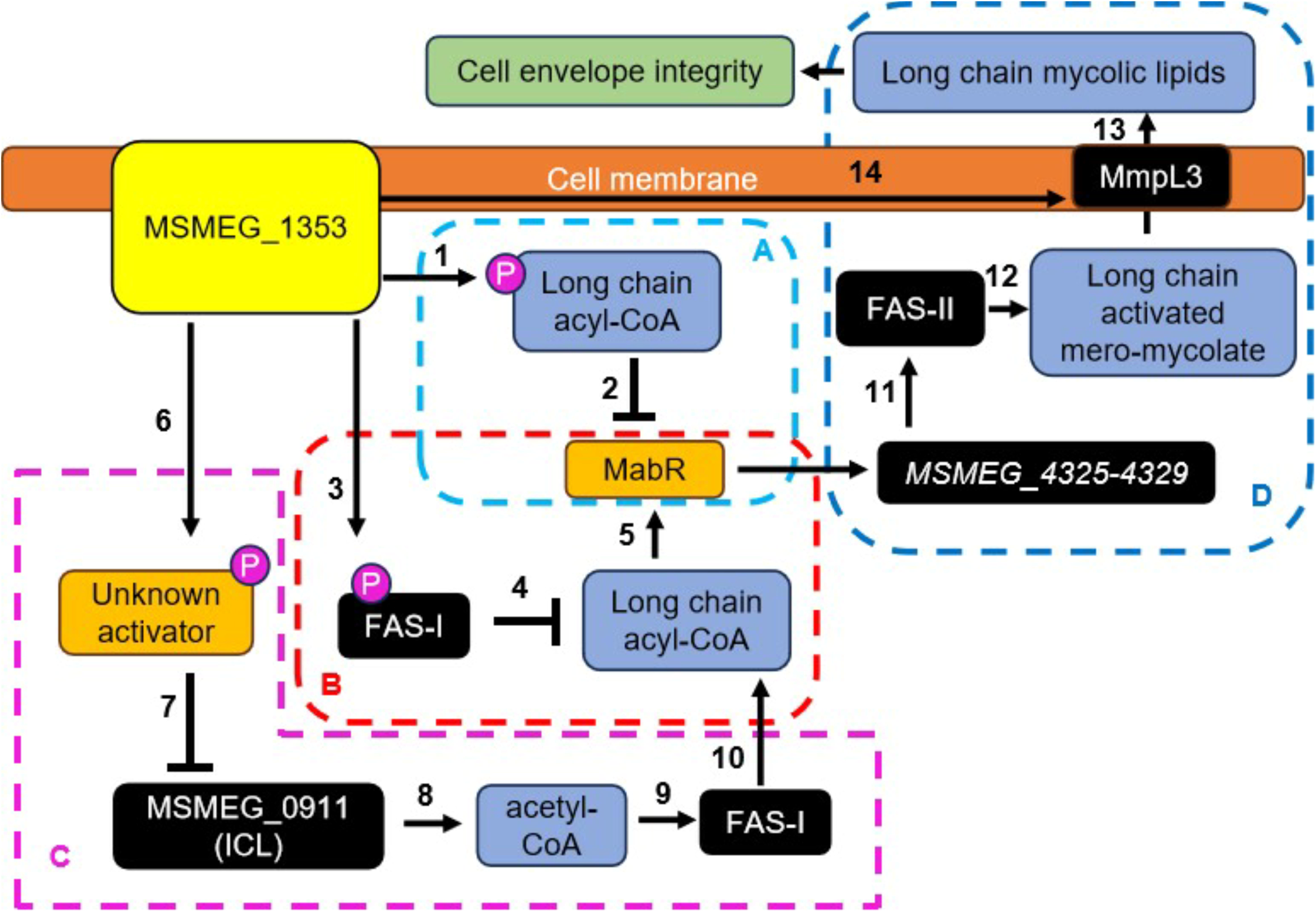
Models for a regulatory role of MSMEG_1353 in mycolic acid synthesis pathways. Based on the experimental results described in this work, we propose different modes of action for MSMEG_1353, acting as a regulator in mycolic acid synthesis pathways. Different pathways enclosed by dashed lines indicated by letters (A, B, C, D) and certain steps are numbered (1-13). **(A)** MSMEG_1353 phosphorylates long-chain acyl-CoAs (step 1) resulting in inhibition of mycolic acid synthesis regulator-MabR (step 2). **(B)** MSMEG_1353 phosphorylates FAS-I proteins (step 3) which reduces the long-chain acyl-CoA production (step 4) and results in less activated MabR (step 5). **(C)** MSMEG_1353 phosphorylates and thereby inhibits an unknown activator for *icl* (step 6), leading to less transcription of *icl* and thus less ICL (step 7), which results in lower amounts of acetyl-CoA as the precursor for FAS-I (steps 8-9). Eventually, this will lead to less long-chain acyl-CoA production (step 10), which negatively affects MabR activity (step 5). **(D)** The *MSMEG_4325-4329* operon activator MabR inhibited via different pathways (A-C) results in less transcription of FAS-II and eventually in lower amounts of long-chain mycolic lipids in the cell envelope (steps 11-13).

Reduced but concentrated corrugation lines on the colony surface of the aTc-induced 1353i strain were seen with the Congo red colony morphology assay. Some *Mtb* proteins involved in modification of cell wall lipids that were transformed and expressed in *MSMEG* were found to result in a wrinkled colony surface, namely Rv3539 (Anand and Kaur, 2023) and Rv0774 (Kumar *et al*., 2017); or in a complete lack of corrugation lines on the colony surface, like with Rv1288 (Maan *et al*., 2018). The decreased aggregation speed and reduced biofilm formation observed in the downregulated strain of *MSMEG_1353* suggest its involvement in the regulation of the mycolic acids biosynthesis pathway, where elongation of mycolic acid can diminish the abilities of *MSMEG* to aggregate and form biofilms (Ojha *et al*., 2005, DePas *et al*., 2019, Gupta *et al*., 2015, Chen *et al*., 2020). The increasing long carbon-chain mycolate by KasA can result in a decreased ability to form biofilms (Ojha *et al*., 2005), similar as the biofilm formation ability decrease seen in this study. An increasing susceptibility to a range of antibiotics and SDS was observed in the aTc-induced 1353i strain. It was noteworthy that SDS, being a detergent that disrupts the cell wall, can be particularly effective against *Mycobacteria* when certain cell wall-associated enzymes are deleted (Meng *et al*., 2017, Ealand *et al*., 2018, Chandra *et al*., 2010). Additionally, this strain exhibited an increased susceptibility to various antibiotics. These characteristics strongly indicate the crucial role played by MSMEG_1353 in maintaining cell wall integrity, potential via affecting biosynthesis of mycolic acids.

An enzymatic cell surface shaving method to determine the proteins exposed on the surface followed by LC-MS/MS analysis showed that MSMEG_1353 is probably located in the cell wall (He and De Buck, 2010). And the predicted 3D structure of MSMEG_1353 by AlphaFold, combined with the conserved amino acids within central regions are potentially part of the active site, which was fitting the ATP binding region of a protein kinase associated with the ABC1_ADCK3-like family prediction from UNIPROT (H121-E448). These may indicate the existence of a cell membrane attachment region, and this region is involved by either or both ends of the protein.

Low MSMEG_1353 results in overexpression of *MSMEG_0911* (*icl*), and the *MSMEG_4325-4329* (*fabD, acpM, kasA, kasB* and *accD6*) operon as detected by MS and RT-qPCR. And genes relative to FasI change were not consistent. These findings were consistent with the upregulation of MSMEG_4327-4328 (KasA and KasB) proteins responsible for carbon-chain elongation in FAS-II process during mycolic acid biosynthesis. Four hypothetical models of how MSMEG_1353 could affect expression of the *FAS-II* operon and subsequent alterations in the carbon-chain length of mycolic acid are displayed in Figure 7 (relevant genes and their homologues are listed in Table S3) and discussed below.

A key player in these hypothetical models is MabR (MSMEG_4324) (Fig. 7A), a regulatory protein in the production of long-chain mycolic acids (Fig. 7D). *Rv2242* (*mabR*) is the homologue of *MSMEG_4324*, which are both shown to be essential genes with a similar function (Salzman *et al*., 2010). Rv2242 (MabR) has been shown to be a transcriptional activator for *fabD, acpM, kasA, kasB* and *accD6* operon (Tsai *et al*., 2017). Interestingly, it seems that long-chain acyl-CoAs (longer than C_18_) can bind MabR (Fig. 7, step 5) and thereby increasing affinity for the operator (Tsai *et al*., 2017). It is possible that when MSMEG_1353 is at a low level, there is an increased availability of long-chain acyl-CoAs for binding to MabR (MSMEG_4324), thereby enhancing the expression of the *MSMEG_4325-4329* (*fabD, acpM, kasA, kasB* and *accD6*) operon and leading to higher FAS-II activity and production of more long-chain mycolic acids (step 5, 11-13).

One explanation for our experimental results could be that MSMEG_1353 phosphorylates long-chain acyl-CoA, as shown in Figure 7A, step 1. This in turn would influence their binding to MabR (MSMEG_4324) (step 2), which results in upregulation of the *MSMEG_4325-4329* operon and more long mycolic acids were transported to the extracellular space (step 11-13). A process that is occurring in parallel or alternatively is shown in area 7B, where MSMEG_1353 phosphorylates FAS-I proteins (step 3), thereby changing their function and eventually resulting in an inhibition of the production of long-chain acyl-CoA (step 4) and subsequent a reduction in the amount of long-chain acyl-CoA that can bind MabR (step 5).

The process shown in area 7C suggests that when MSMEG_1353 is present at low levels, MSMEG_0911 (ICL) could function to maintain acetyl-CoA at an optimal level for further mycolic acid biosynthesis (step 6-10). This aligns with our observation that expression of MSMEG_0911 (ICL) negatively correlates with MSMEG_1353. During hypoxic conditions, the generation of pyruvate from glycolysis was reduced, and MSMEG_0911 (ICL) as the first enzyme in the glyoxylate cycle leads to a bypass of the CO_2_-generating steps in the tricarboxylic acid cycle. The glyoxylate cycle works in concert with the tricarboxylic acid cycle to maintain oxaloacetate levels and sustain acetyl-CoA production (Gould *et al*., 2006). The acetyl-CoA is an important precursor for mycolic acid biosynthesis and mycolic acids is important to maintaining cell envelope integrity, and the cell envelope integrity was disrupted when *MSMEG_1353* was downregulated, and mycolic acid elongation enzymes overexpressed in aTc-induced 1353i strain. These findings suggest that MSMEG_0911 (ICL) may be indirectly inhibited via MSMEG_1353.

A final hypothesis for some of the observed results is indicated by arrow number 14 where MSMEG_1353 is depicted to inhibit MmpL3 via an unknown process to reduce mycolic acid transportation. MmpL3 is accountable for transporting a type of mycolic acid from the cytoplasm to the inner membrane of the *Mtb*, which is regarded as a promiscuous drug target (Imran *et al*., 2022).

Taken together, the findings described in this study have led to the postulation of several models that each provide a possible explanation for how it could be involved in mycolic acid biosynthesis. Which of these models is a true reflection of reality remains to be determined experimentally in the future.

## Supporting information

Graphical abstract

## Author contributions

**Ziwen Xie:** Conceptualization, Methodology, Validation, Formal analysis, Investigation, Data Curation, Writing - Original Draft, Writing - Review & Editing, Visualization, Project administration **Trillion Surya Lioe:** Methodology, Investigation **Abhipsa Sahu:** Methodology **Jiahao Cui:** Methodology, Investigation, Data Curation **David Ruiz-Carrillo:** Conceptualization, Supervision, Funding acquisition **Tatsuhiko Kadowaki:** Supervision, Funding acquisition **Boris Tefsen:** Conceptualization, Methodology, Resources, Writing - Review & Editing, Supervision, Project administration, Funding acquisition

## Declaration of Competing Interest

None of the authors has any financial or personal relationships that could inappropriately influence or bias the content of the paper.

## Data Availability

Data will be made available on request.

## Acknowledgements

This project was financially supported by the Xi’an Jiaotong Liverpool University doctoral scholarship (PGRS1906022). We sincerely thank Mal Horsburgh for co-supervising this project.

## Notes

### Competing Interest Statement

The authors have declared no competing interest.

