## supplemental figure legend for "Linking MSMEG_1353 to lipid metabolism and envelope integrity in *Mycolicibacterium smegmatis*"

**Fig. S1. Alignment of MSMEG_1353 and Rv0647c.**

The alignment of both MSMEG_1353 and Rv0647c was created using Clustal Omega (Madeira *et al.*, 2019) and ESPript 3.0 (Robert and Gouet, 2014). Red represents conserved amino acids, while yellow represents similar amino acids.

**Fig. S2. Bacterial dimensions examined by light microscope of strains having differential *MSMEG_1353* expression.**

Cell length and width of each strain were measured for 90 individual bacteria from three replicate populations, using ImageJ, with data analysis by GraphPad. **(A)** Length and **(C)** width values of cells under light microscope. **(B)** Cell samples of strains grown in the presence or absence of aTc under light microscope at 630× magnification. Results are the mean of three independent biological experiments performed in triplicate each, and the error bars represent ± standard deviation. Asterisks indicate significant differences in an unpaired t test (*** $p < 0.001$; **** $p < 0.0001$).

**Fig. S3. Ratio of cell length to width under SEM of strains with differential *MSMEG_1353* expression.**

Data for cell length (L) and width (W) of each strain were collected from SEM photos and the ratio was calculated as L/W. Results are the mean of three independent biological experiments performed in triplicate each, and the error bars represent ± standard deviation. Asterisks indicate significant differences in an unpaired t test (* $p < 0.05$).

**Fig. S4. Calculated cell volume under SEM overview of strains with differential *MSMEG_1353* expression.**

Data for cell length (L) and width (W) of each strain were collected from SEM photos and cell volume value was calculated by $V = \frac{\pi W^2}{4(L - \frac{W}{3})}$. Results are the mean of three independent biological experiments performed in triplicate each, and the error bars represent ± standard deviation. Asterisks indicate significant differences in an unpaired t test (*** $p < 0.001$; **** $p < 0.0001$).

**Fig. S5. Aggregation index at t = 5 min of strains with differential *MSMEG_1353* expression.**

Aggregation index at t = 5 min of strains grown in 7H9 medium without Tween-80 for 2 days in the presence or absence of aTc. Results are the mean of five independent biological experiments performed in triplicate each, and the error bars represent ± standard deviation. Asterisks indicate significant differences in an unpaired t test (* $p < 0.05$; ** $p < 0.01$).

**Fig. S6. Inhibition of strains with differential *MSMEG_1353* expression in an SDS diffusion assay.**

Representative example of strains grown in the presence or absence of aTc on 7H10 plates for discs containing different concentrations of SDS. Three independent experiments were performed, with each strain tested in triplicate.

**Fig. S7. Inhibition of strains with differential *MSMEG_1353* expression on agar plates containing SDS.**

Representative example of strains grown in the presence or absence of aTc on the 7H10 agar plates with different concentrations of SDS. Three independent experiments were performed, with each strain tested in triplicate.

**Fig. S8. Alignment of MSMEG_1353 homologues.**

The secondary structures of partly full-length MSMEG_1353 (47F-324L) were illustrated at the top of the alignment, with the model based on the 5yjz structure. Continuous nonconservative protein fragments at both ends of full-length MSMEG_1353 were not shown. Regions that were completely conserved were depicted in red, while highly conserved regions were shown in yellow. Figure obtained from ESPript (https://espript.ibcp.fr/ESPript/ESPript/).

**Fig. S9. Predicted 3D structure of full-length MSMEG_1353 based on AlphaFold.**

The black arrow displays the N-terminus of full-length MSMEG_1353. The structure is colored by pLDDT score. Residues in the 3D viewer were color-coded using confidence bands (dark blue > 90, light blue: 70 to 90, yellow: 50 to 70, orange < 50). Figures were generated with PyMOL version 2.3.2.

**Fig. S10. Volcano plot of *MSMEG* proteins identified by MS.**

Proteins exhibiting a positive $\log_2$ fold change were found to be upregulated in the proteome, while those with a negative fold change indicated protein expression inhibition compared to the wt strain. A greater magnitude of fold change corresponded to a higher degree of overexpression or inhibition. The two vertical dotted lines indicate a $\pm$ 1.5-fold change. The Y-axis represents the *p*-value, where values above the horizontal dotted line indicate a *p*-value lower than 0.05. **(A-C)** depict volcano plots for the aTc-induced 1353i strains from three independent experiments, while **(D-F)** represent volcano plots for the 1353up strains from three independent experiments. Points colored red denote proteins identified as being inhibited compared to the wt strain. Points colored blue signify proteins identified as being overexpressed compared to the wt strain. Points colored grey either have a *p*-value higher than 0.05 or exhibit a fold change lower than 1.5.
