## supplemental figure 1 for "Linking MSMEG_1353 to lipid metabolism and envelope integrity in *Mycolicibacterium smegmatis*"

```
               1        10        20        30        40        50        60
Mtb_Rv0647c    MRAEIGPDFRPHYTFGDAYPASERAHVNWELSAPVWHTAQMGSTTHREVAKLDRVPLPVE
MSMEG_1353     ....................................MSTTPAKTKHREVARLDRVPLPAE

                        70        80        90       100       110       120
Mtb_Rv0647c    AARVAATGWQVTRTAVRFIGRLPRKGPWQQKVIKELPQTFADLGPTYVKFGQIIASSPGA
MSMEG_1353     AARIGATGWQITRTGARVAANIFGRGSLQQKIVKQVPKTFSDLGPTYVKFGQIIASSPGA

                       130       140       150       160       170       180
Mtb_Rv0647c    FGESLSREFRGLLDRVPPAKTDEVHKLFVEELGDEPARLFASFEEEPFASASIAQVHYAT
MSMEG_1353     FGEPLSREFRGLLDRVPPANPDEVQKLIREELGDDPHKLFKTFDDTPFASASIAQVHYAT

                       190       200       210       220       230       240
Mtb_Rv0647c    LRSGEEVVVKIQRPGIRRRVAADLQILKRFAQTVELAKLGRRLSAQDVVADFADNLAEEL
MSMEG_1353     LHSGEEVVVKIQRPGIRRRVAADLQILKRGARLVEFAKLGQRLSAQDVVADFADNLAEEL

                       250       260       270       280       290       300
Mtb_Rv0647c    DFRLEAQSMEAWVSHLHASPLGKNIRVPQVHWDFTTERVLTMERVHGIRIDNAAAIRKAG
MSMEG_1353     DFRLEAQSMDAWVAHMHASPLGANIRVPTVYWDLTSERVLTMERITGVRIDDVAAIRKKG

                       310       320       330       340       350       360
Mtb_Rv0647c    FDGVELVKALLFSVFEGGLRHGLFHGDLHAGNLYVDEAGRIVFFDFGIMGRIDPRTRWLL
MSMEG_1353     FDGTELVKALLFSVFEGGLRHGLFHGDLHAGNLYVDDDGKIVFFDFGIMGRIDPRTRWLL

                       370       380       390       400       410       420
Mtb_Rv0647c    RELVYALLVKKDHAAAGKIVVLMGAVGTMKPETQAAKDLERFATPLTMQSLGDMSYADIG
MSMEG_1353     RELVHALLVKKDHAAAGKIVVLMGAVGTVKPEGQAAKDLEAFATPLTMKSLGDMSYAEIG

                       430       440       450       460       470       480
Mtb_Rv0647c    RQLSALADAYDVKLPRELVLIGKQFLYVERYMKLLAPRWQMMSDPQLTGYFANFMVEVSR
MSMEG_1353     RQLSALADAYDVKLPRELVLIGKQFLYVERYMKLLAPKWQMMSDPQLTGYFANFMVEVSR

Mtb_Rv0647c    EHQSDIEV.....................
MSMEG_1353     EHKEIDDVDEIPDIPETGTETQAPSGDKA
```
