## Supplementary figures and images for "Linking MSMEG_1353 to lipid metabolism and envelope integrity in *Mycolicibacterium smegmatis*"

### Graphical abstract

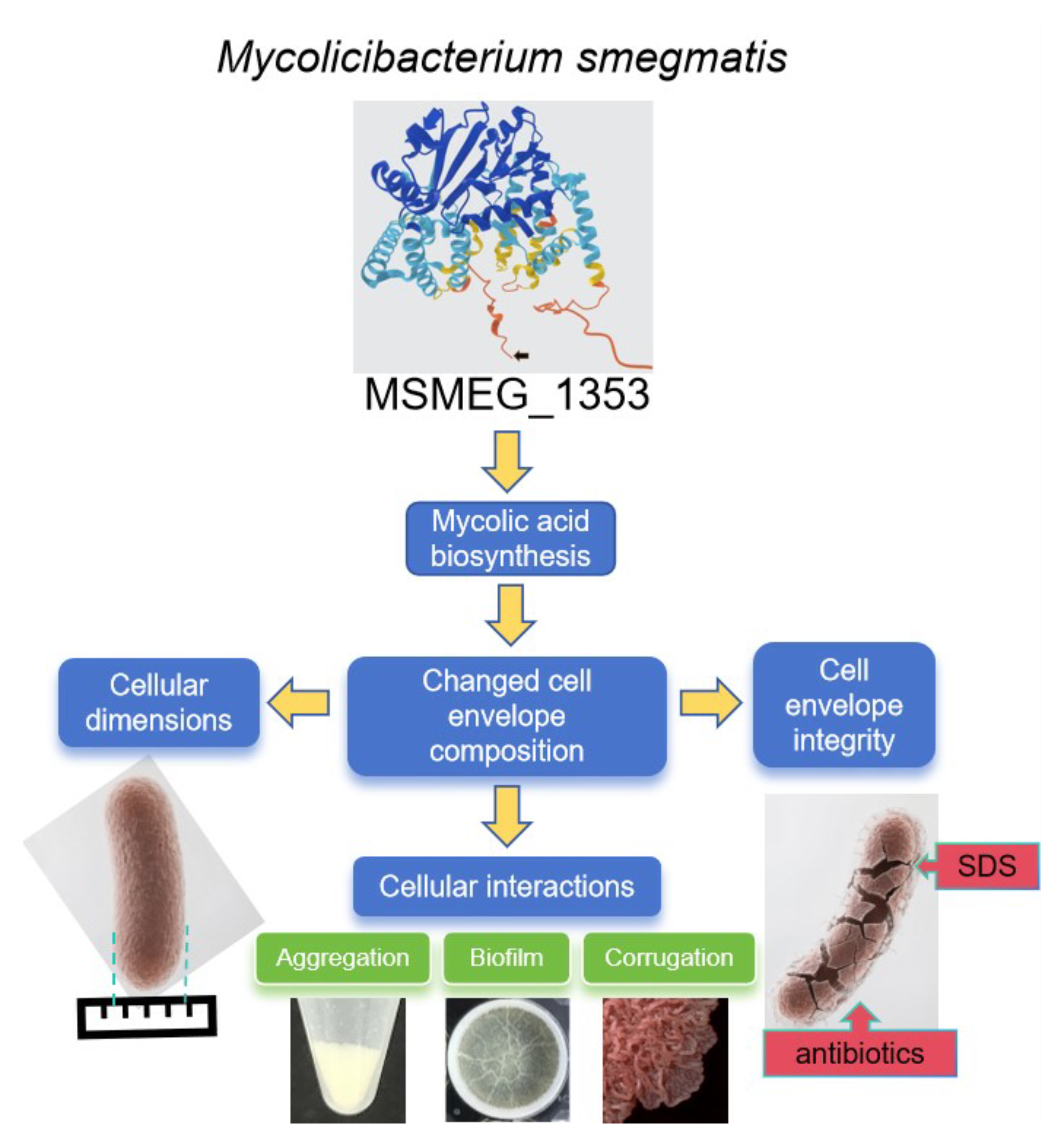

### supplemental figure 2

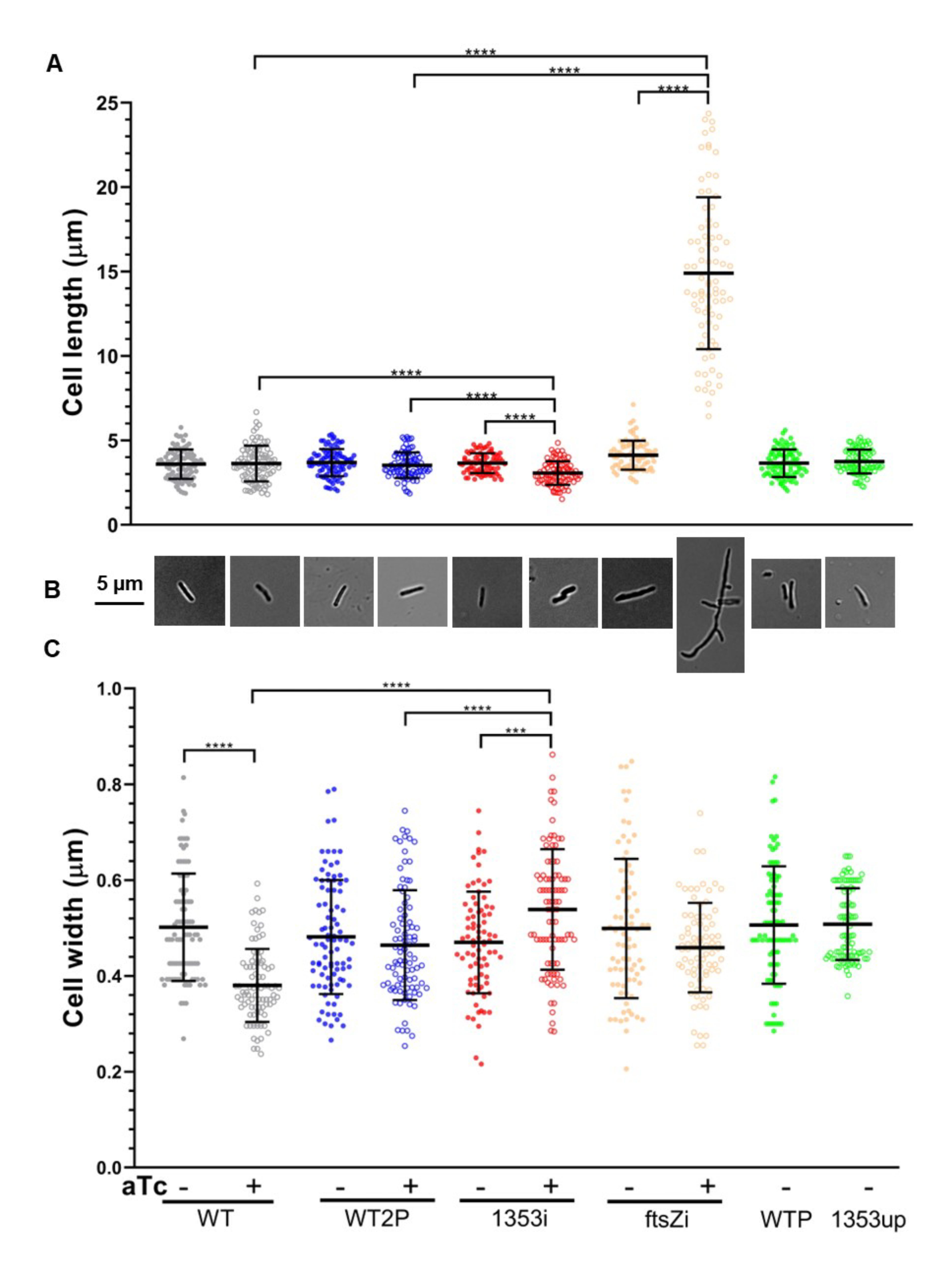

### supplemental figure 3

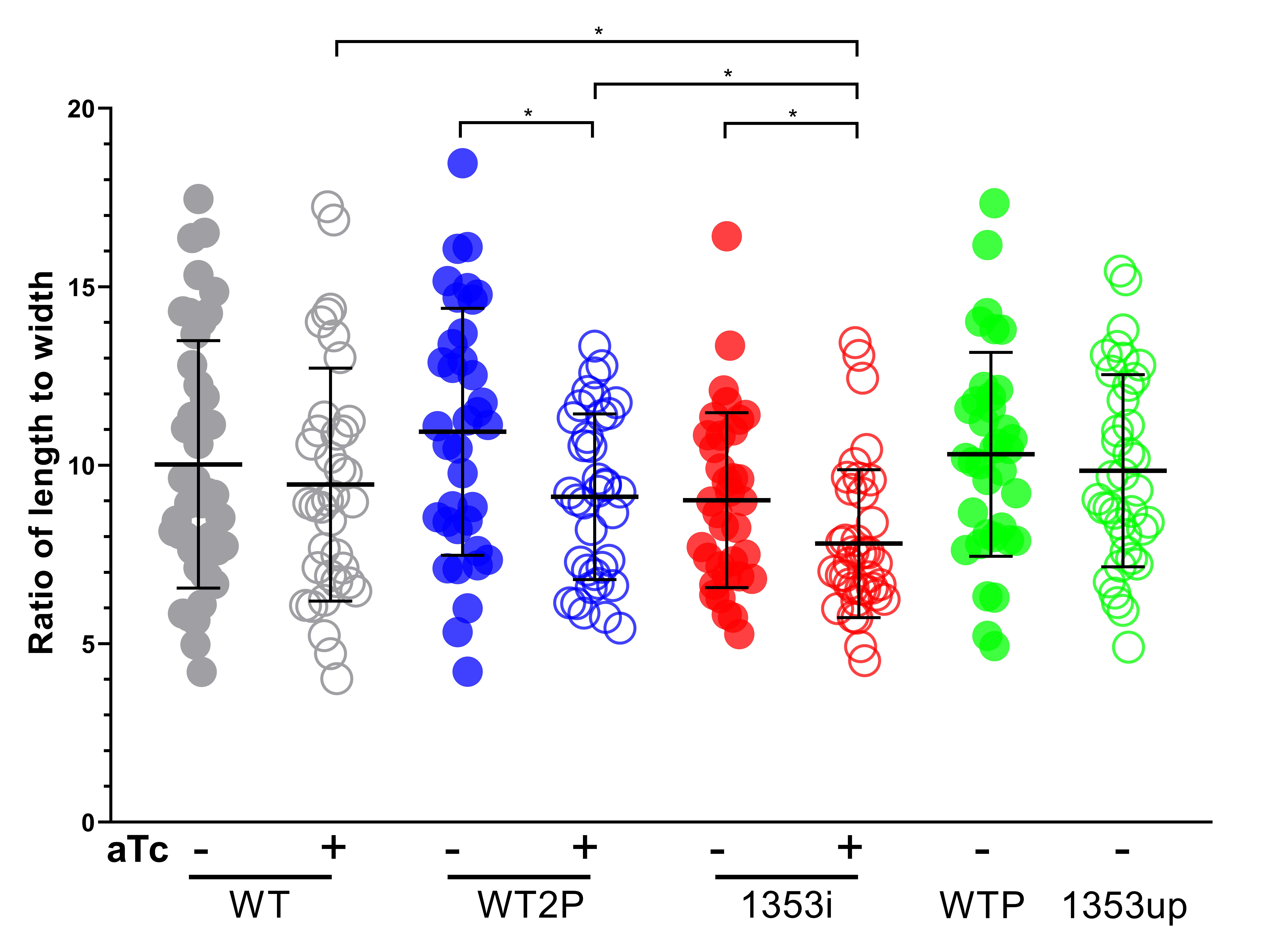

### supplemental figure 4

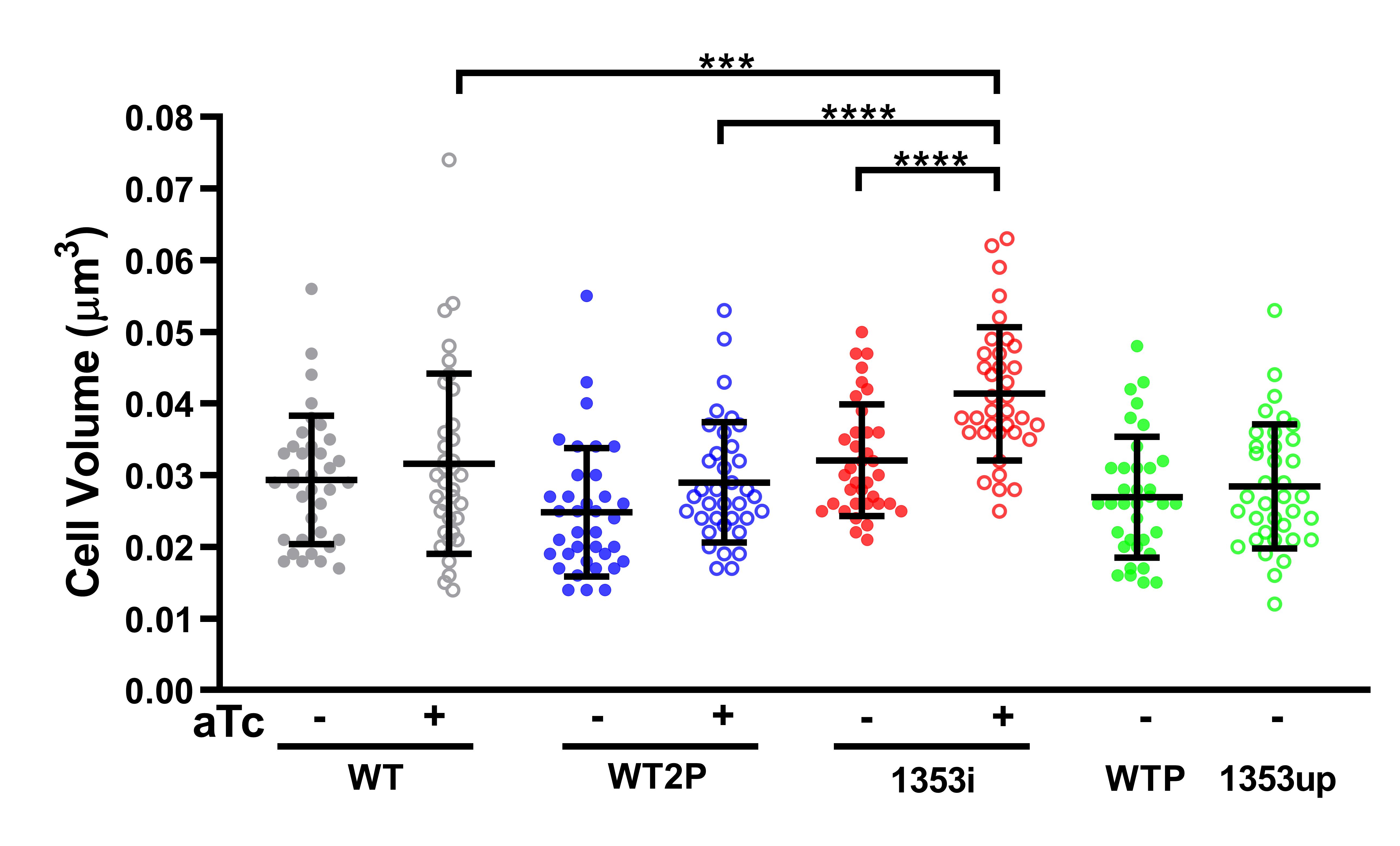

### supplemental figure 5

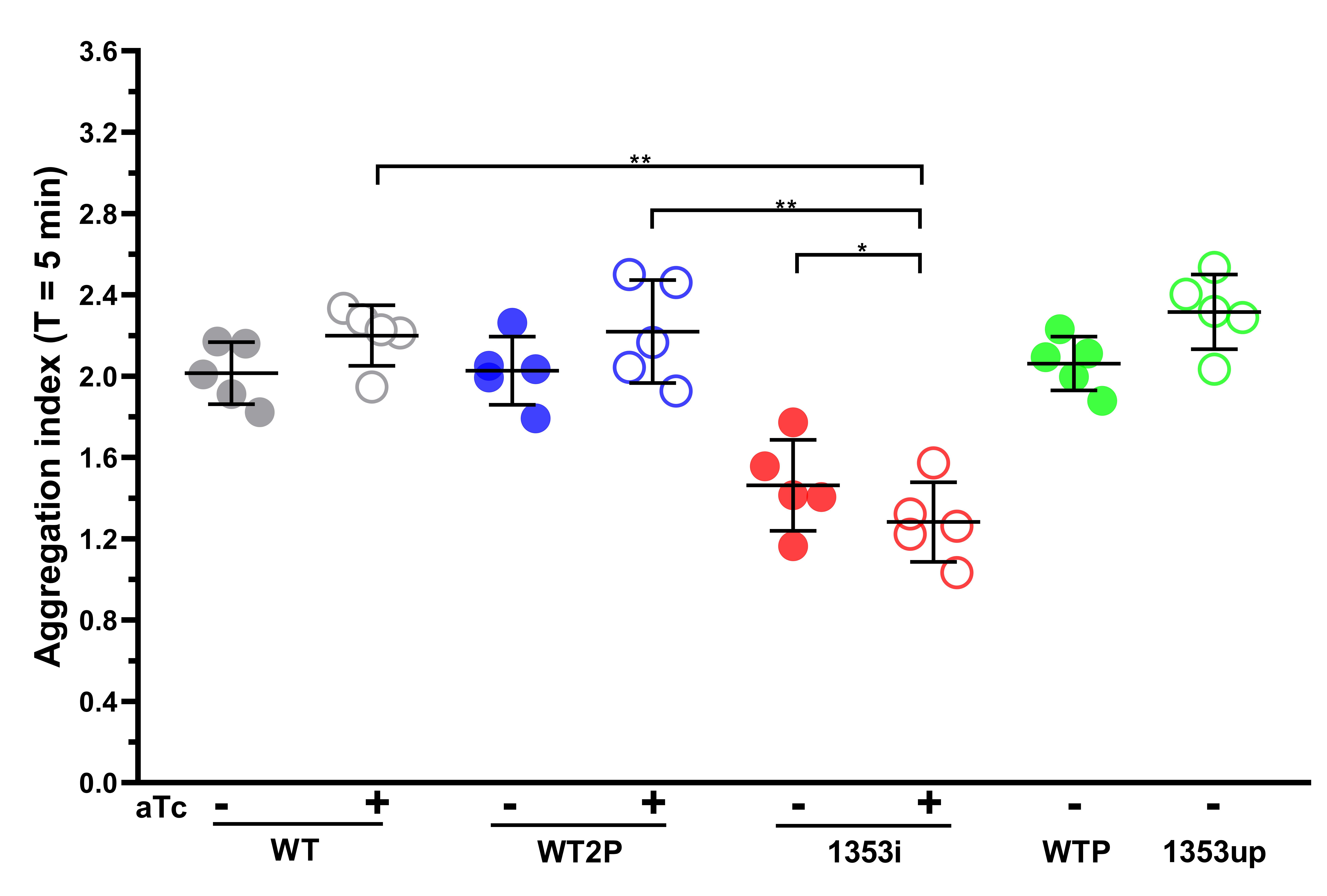

### supplemental figure 6

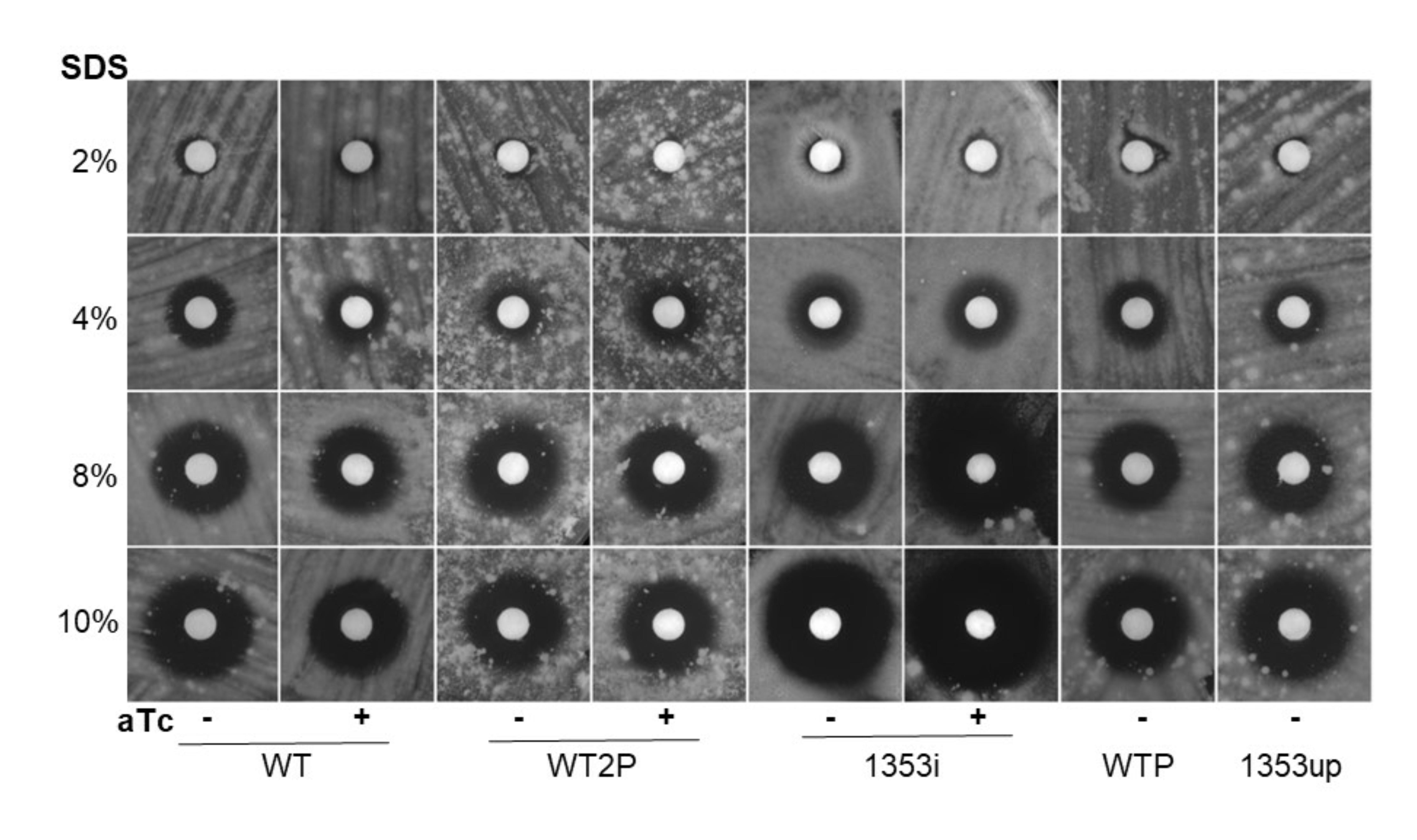

### supplemental figure 7

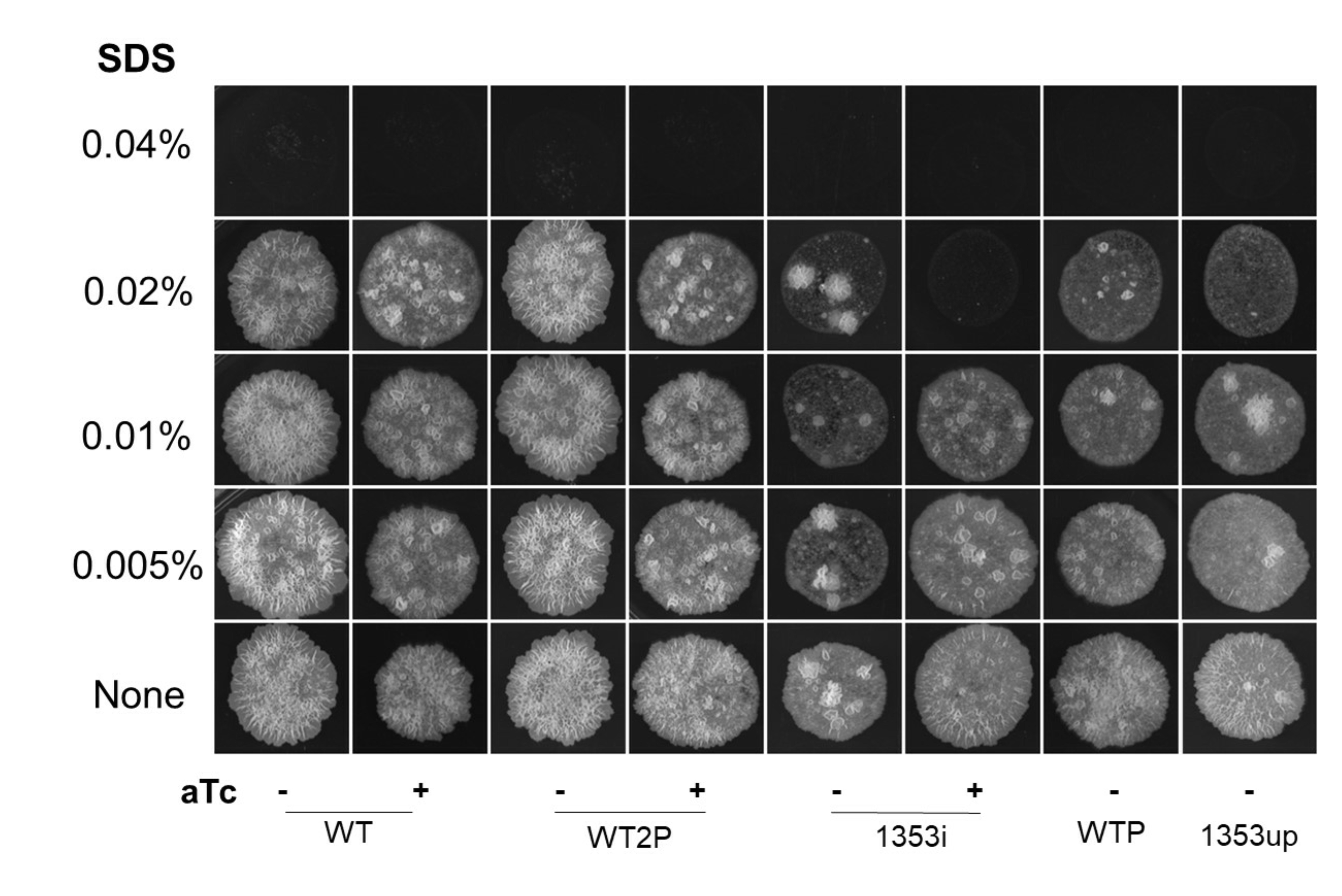

### supplemental figure 8

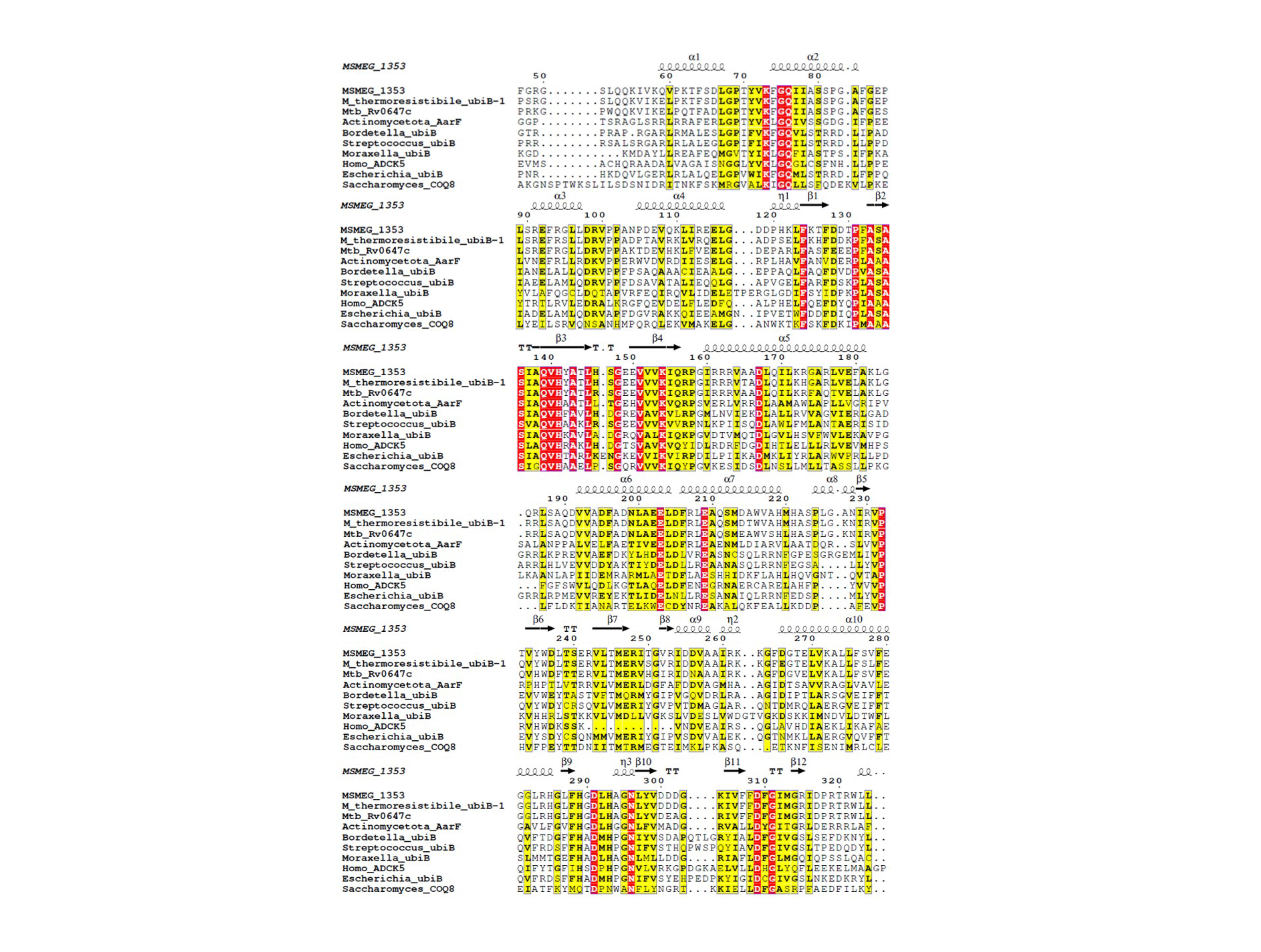

### supplemental figure 9

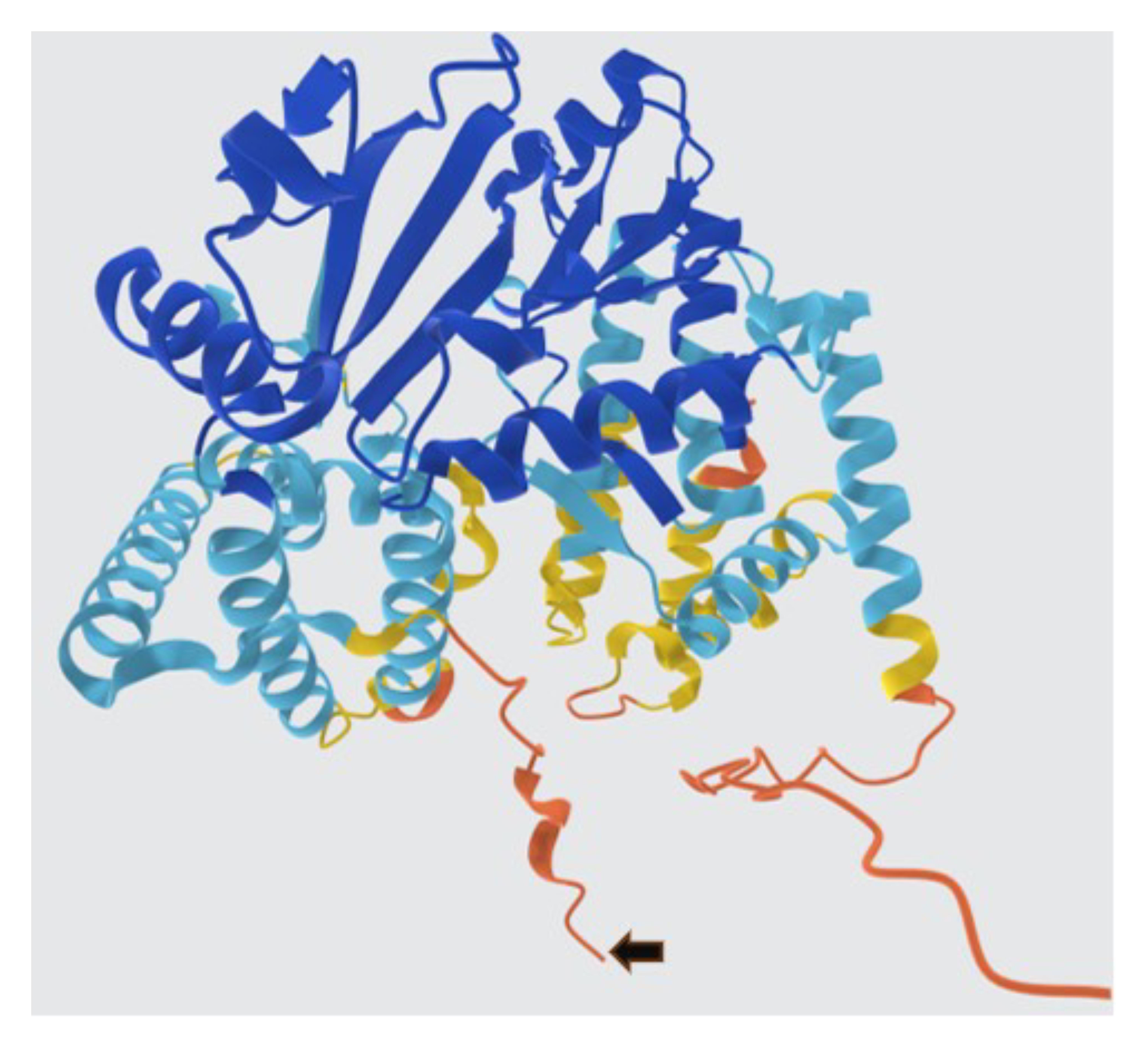

### supplemental figure 10

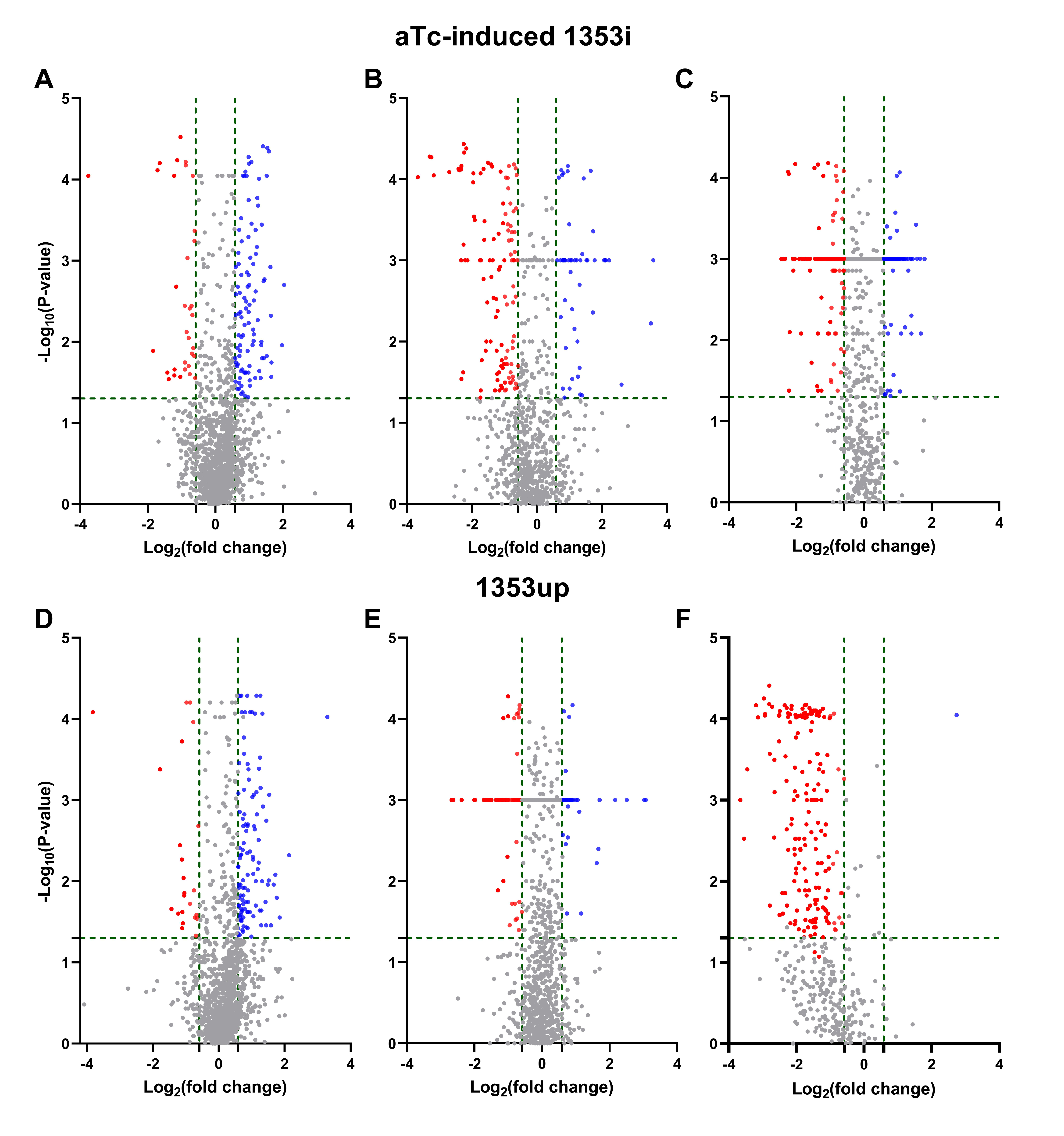
