## supplemental table 1-3 for "Linking MSMEG_1353 to lipid metabolism and envelope integrity in *Mycolicibacterium smegmatis*"

**Table S1**

Strains and plasmids used in this study.

| Strain or plasmid | Description |
|---|---|
| ***E. coli*** | |
| DH5α | Strain used for general cloning procedures |
| ***MSMEG*** | |
| $mc^2155$ (WT) | Wild-type laboratory strain; Genomic DNA used as polymerase chain reaction (PCR) template |
| WTP | $Kan^R$; $mc^2155$ carrying pMV261; used as negative control |
| 1353up | $Kan^R$; $mc^2155$ carrying pMV261-1353; *MSMEG_1353* overexpression strain |
| WT2P | Hygromycin $(Hyg)^R$ and $Kan^R$; $mc^2155$ carrying pRH2502 and pRH2521; used as negative control |
| 1353i | $Hyg^R$ and $Kan^R$; $mc^2155$ carrying pRH2502 and pRH2521-1353; inducible *MSMEG_1353* knockdown strain |
| ftsZi | $Hyg^R$ and $Kan^R$; $mc^2155$ carrying pRH2502 and pRH2521-ftsZ; inducible *ftsZ* knockdown strain |
| **Plasmids** | |
| pMV261 | $Kan^R$; *Mycobacterium bovis* hsp60 used as promoter; used as carrier of *MSMEG_1353* gene insertion |
| pMV261-1353 | $Kan^R$; Plasmid carries *MSMEG_1353* gene; for *MSMEG_1353* gene overexpression |
| pRH2502 | $Kan^R$; Expression of dCas9 (D10A H840A) from a TetR-regulated uvtetO promoter; used for CRISPRi |
| pRH2521 | $Hyg^R$; Vector for sgRNA insertion expressed from a TetR-regulated smyc promoter (Pmyc1tetO); used for CRISPRi |
| pRH2521-ftsZ | $Hyg^R$; carrying *ftsZ* for conditional knockdown |
| pRH2521-1353 | $Hyg^R$; carrying *MSMEG_1353* for conditional knockdown |

$^R$ resistance

**Table S2**

Homologues of full-length MSMEG_1353 in selected organisms.

| Species | Protein name | Percentage Identity | Accession code |
|---|---|---|---|
| *MSMEG* | MSMEG_1353 | 100% | WP_011727618.1 |
| *Mycolicibacterium thermoresistibile* | UbiB | 87.11 % | WP_003926000.1 |
| *Mtb* | Rv0647c | 84.09 % | WP_003403317.1 |
| *Actinomycetota bacterium* | AarF | 37.22 % | TML89699.1 |
| *Bordetella pertussis* | UbiB | 32.05 % | WP_080485500.1 |
| *Streptococcus pneumoniae* | UbiB | 32.02 % | CJK64186.1 |
| *Moraxella catarrhalis* | UbiB | 30.59 % | WP_152699611.1 |
| *Homo sapiens* | ADCK5 | 30.28 % | EAW82111.1 |
| *Escherichia coli* | UbiB | 30.17 % | MWO99691.1 |
| *Saccharomyces cerevisiae* | COQ8 | 29.01 % | AJS05093.1 |

**Table S3**

Enzymes from mycolic acid biosynthesis pathway detected by MS.

| Pathway | ID in *Mtb* | ID in *MSMEG* | Enzyme | Fold change direction | |
|---|---|---|---|---|---|
| | | | | **1353i+aTc** | **1353+** |
| FAS-I | Rv2524c | MSMEG_4757 | Fatty acid synthetase-I | 2↓ | 3↑ |
| FAS-I→FAS-II | Rv2243 | MSMEG_4325 (FabD) | Malonyl-CoA:ACP transacylase | 1↑ | 1↓ |
| | Rv2244 | MSMEG_4326 (AcpM) | acyl carrier protein | 1↑ | 1↓ |
| FAS-II | Rv0635 | MSMEG_1340 (HadA) | (3R)-hydroxyacyl-ACP dehydratase subunit HadA | 1↓ | Nd |
| | Rv0637 | MSMEG_1342 (HadC) | (3R)-hydroxyacyl-ACP dehydratase subunit HadC | 1↓ | 1↑ |
| | Rv2245 | MSMEG_4327 (KasA) | beta_-Ketoacyl-ACP synthase | 1↑ | 1↓ |
| | Rv2246 | MSMEG_4328 (KasB) | beta_-Ketoacyl-ACP synthase | 3↑ | 2↓ |
| | Rv1141c | MSMEG_5185 | enoyl-CoA hydratase PaaB | 1- | 1- |
| | Rv1484 | MSMEG_3151 (InhA) | 2-*trans*-Enoyl-ACP reductase | 1- | 1↓ |

Downregulated proteins are indicated by down arrows; upregulated proteins are indicated by up arrows, the number before the arrows indicate the times this protein was detected among the three independent experiments for that strain. Nd not detected.
